# LCK deficiency in CD8 T cells leads to reduced proliferation and increased effector T-cell formation in mice

**DOI:** 10.1101/2025.08.28.672860

**Authors:** Valeria Uleri, Vojtech Racek, Marta Popovic, Anna Morales Mendez, Veronika Niederlova, Arina Andreyeva, Juraj Michalik, Nadine M. Woessner, Michaela Krupkova, Radislav Sedlacek, Susana Minguet, Ondrej Stepanek

## Abstract

LCK is an SRC-family kinase that mediates the initial steps in T-cell antigen receptor signaling and governs positive and negative selection during thymocyte development. While its developmental role is well established, its functions in peripheral T-cell responses remain poorly defined. Here, we investigated the responses of wild-type and LCK-deficient TCR-transgenic OT-I T cells across two infection models and an autoimmune diabetes model. LCK-deficient T cells exhibited reduced antigen-induced proliferation but, paradoxically, displayed enhanced effector differentiation in vivo. This phenotype likely reflects dysregulation of specific TCR signaling pathways, as LCK was more critical for ERK and NFAT activation than for NFκB, AP-1, or AKT/mTOR signaling. T cells deficient in a related kinase FYN also showed a slight increase in effector cell formation, suggesting that effector differentiation is regulated by their combined activity rather than distinct non-redundant roles. Our results reveal that LCK has two intrinsic roles in T-cell responses – promoting proliferation while restraining effector differentiation. These findings provide new insight into the molecular mechanisms of T-cell activation in vivo with implications for understanding the pathophysiology of LCK deficiency in humans and optimizing adoptive T-cell therapies.

## Introduction

Antigen receptor signaling is initiated by the phosphorylation of immunoreceptor tyrosine-based activation motifs (ITAM) by SRC-family tyrosine kinases (SFK). LCK is considered as the major SFK (*1*) in T-cell receptor (TCR) signaling. It phosphorylates the ITAMs in the TCR-associated CD3 and ζ chains, as well as the kinase ZAP-70, which is subsequently recruited to doubly phosphorylated ITAMs. LCK is specifically expressed in T cells and is the only SFK interacting with the T-cell co-receptors CD4 and CD8 (*2*). Whereas the CD4-LCK interaction is important for the development and responses of CD4^+^ T cells, the CD8-LCK interaction plays a relatively minor role in CD8^+^ T cells (*3*).

LCK-deficient thymocytes exhibit defective pre-TCR signaling-mediated β-selection and TCR signaling-mediated positive selection leading to partial blocks in thymocyte development at the DN3 and DP stages, respectively in mice (*3-6*). These developmental arrests result in low peripheral T-cell counts (*3-6*). Complete or partial LCK deficiency in humans is classified as an autosomal recessive disorder characterized by low peripheral T-cell numbers, intrinsic T-cell unresponsiveness to TCR activation, recurrent infections, and autoimmune symptoms (*7-9*). CD4^+^ T cells are generally more affected than CD8^+^ T cells in these patients.

Another SFK member expressed in T cells is FYN. In contrast to LCK, FYN-deficient mice show essentially normal development and antigenic responses of conventional T-cells with only minor differences in comparison to wild-type (WT) mice (*10, 11*). In the absence of LCK, FYN supports partially the thymocyte maturation as suggested by the complete thymocyte developmental arrest in *Lck*^-/-^ *Fyn*^-/-^ double knock-out (KO) mice (*4, 5, 12*).

The roles of LCK and FYN in the effector responses of mature T cells are not fully understood. One reason is the difficulty of distinguishing the effects of LCK in peripheral T cells from its earlier role in thymic selection, since LCK deficiency alters T-cell maturation in both cell numbers and TCR repertoire. A few studies have suggested that LCK is not essential for antigenic responses of memory T cells (*13*) or the function of chimeric antigen receptors (CARs) expressed in human T cells (*14*). Moreover, murine LCK-deficient T cells exhibited reduced, but not completely blocked TCR responses in vitro (*15, 16*). The magnitude of the defect in LCK-deficient T cells in vitro varied considerably depending on the assay, readout, and study (*15-17*). Thus, the impact of LCK deficiency on intrinsic T-cell effector responses in vivo remains largely unknown.

In this study, we investigated the role of LCK and FYN in peripheral CD8^+^ T-cell responses using adoptive transfers of LCK or FYN-deficient monoclonal T cells into polyclonal hosts. This strategy allowed us to bypass the quantitative and qualitative thymic selection defects coupled with SFKs deficiency. Upon in vivo stimulation, LCK-deficient CD8^+^ T cells showed weaker expansion coupled to a preferential differentiation to effector T cells. Remarkably, LCK was largely dispensable for the in vivo induction of experimental diabetes by CD8^+^ T cells. These results uncover a previously unknown context-dependent interplay among SFKs in T cells. While FYN plays only a minor role in LCK-sufficient T cells, it can partially compensate for the absence of LCK in *Lck*^-/-^ T cells and is required for the development of CD8^+^ T cells in mice with disrupted CD8-LCK interaction.

## Results

### LCK-deficient T cells respond to cognate pathogens in vivo

To analyze the role of LCK in mature CD8^+^ T cells, we used *Lck* WT (henceforth only WT), *Lck*^-/-^, and *Lck*^CA/CA^ (LCK is unable to bind to CD4 and CD8 co-receptors in these mice) (*3*) on the background of TCR transgenic OT-I *Rag2*^-/-^ (henceforth OT-I) mice. The fixed OT-I TCR was used to avoid alterations in the TCR repertoire observed in *Lck*^-/-^ mice due to the impaired positive and negative selection in the thymus, and to obtain a monoclonal population of CD8^+^ T cells specific for ovalbumin (OVA)-derived peptide. To overcome the fact that peripheral T-cell numbers were reduced in the *Lck*^-/-^ OT-I mice compared to their *Lck* WT counterparts (Supplemental Fig. 1A-B), we transferred equal numbers of WT, *Lck*^-/-^, or *Lck*^CA/CA^ OT-I T cells into polyclonal host mice to evaluate their antigenic responses on a per-cell basis.

First, we adoptively transferred numbers of unsorted lymphocytes from WT or *Lck*^*−/−*^ OT-I *Rag2*^*−/−*^ mice corresponding to 1×10^4^ OT-I CD8^+^ T cells into congenic Ly5.1 host mice and assessed their response to a subsequent infection with *Listeria monocytogenes* expressing their cognate antigen OVA (LM-OVA) (*18*) (Fig. 1A). Given the established central role of LCK in T-cell activation, we expected LCK-deficient cells to be largely unresponsive. However, *Lck*^-/-^ OT-I T cells retained a substantial capacity to expand upon LM-OVA infection on day six post-infection, revealing that intrinsic in vivo T-cell responses occur in the absence of canonical LCK signaling (Fig. 1B, Supplemental Fig. 1C). Moreover, the phenotypic analysis of these cells revealed that *Lck*^-/-^ OT-I T cells upregulated KLRG1, a marker of short-lived effector T cells, as well as CD62L, a marker of central memory T cells (Fig. 1B-C). To avoid any potential artifacts caused by the transfer of non-T cells, we reproduced the experiment using 1×10^4^ OT-I T cells FACS-purified as CD5^+^ CD8^+^ double-positive lymphocytes for the adoptive transfer (Supplemental Fig. 2A-B). We also included *Lck*^CA/CA^ OT-I cells for comparison. Again, we observed a modest expansion and a significant upregulation of KLRG1, and to a lesser extent, other effector markers such as KLRK1 and CX3CR1, in *Lck*^-/-^ OT-I cells compared to WT counterparts (Fig. 1D, Supplemental Fig. 2C). *Lck*^CA/CA^ OT-I cells showed a comparable response to WT OT-I cells as previously described (*3, 19*).

**Figure 1.**
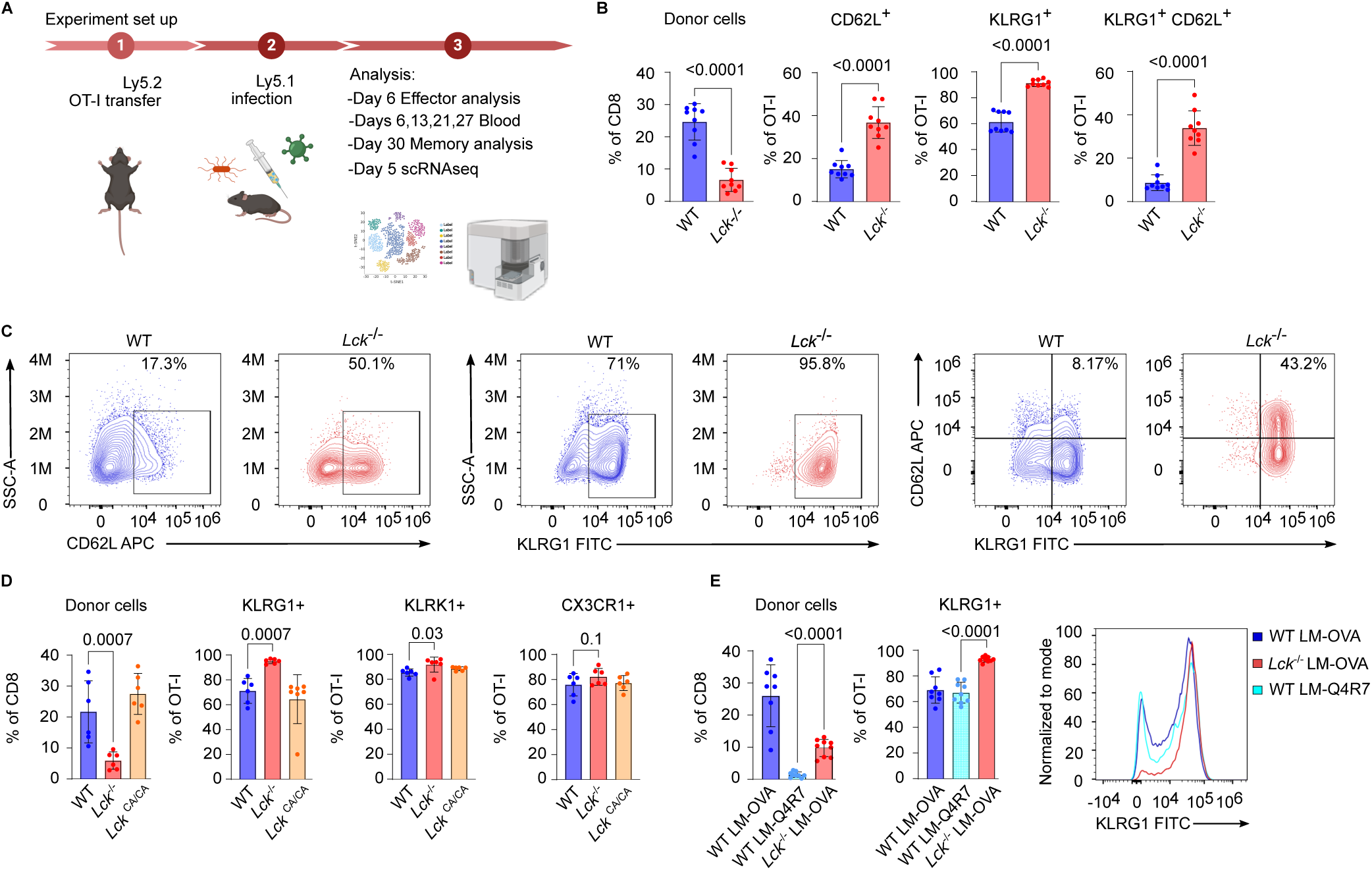
LCK-deficient CD8^+^ T cells respond to cognate bacterial infection in vivo. (A) Schematic representation of infection experiments performed in this study. (B-E) 1×10^4^ T cells from WT, *Lck*^CA/CA^, or *Lck*^-/-^ OT-I *Rag2*^-/-^ mice were adoptively transferred to Ly5.1 host mice that were subsequently infected with LM-OVA (B-E) or LM-Q4R7 (E). Splenocytes were analyzed by flow cytometry on day six post-infection. The adoptively transferred OT-I T cells were counted as CD8α^+^ TCRβ^+^ and injected alongside non-T cells isolated from the lymph node and spleens from indicated mice (B-C) or FACS-sorted as CD8α^+^ CD5^+^ (D-E) (B-C) Aggregate results (B) and representative contour plots (C) show the frequency of transferred OT-I T cells and their CD62L and KLRG1 expression. n = 9 mice per group in 4 independent experiments. (D) Aggregate results show the frequency of transferred OT-I T cells and their KLRG1, KLRK1, and CX3CR1 expression. n = 6 mice per group in 2 independent experiments. (E) Aggregate results and a representative histogram of KLRG1 expression. n =8 mice for WT/LM-OVA, n = 9 mice for WT/LM-Q4R7 and *Lck*^-/-^/LM-OVA. Mean±SEM is shown. Statistical significance was calculated using two-tailed Mann-Whitney test.

The bias toward effector differentiation in *Lck*^-/-^ OT-I T cells was in apparent contrast to a previous report showing that suboptimal TCR activation by low-affinity antigens favors the memory-precursor phenotype in CD8^+^ T cells (*20*). To test this directly, we compared *Lck*^-/-^ OT-I cells responding to high affinity LM-OVA with WT OT-I cells responding to LM expressing a lower affinity variant of OVA (LM-Q4R7). Although the expansion of WT OT-I responding to LM-Q4R7 was much reduced compared to LM-OVA, they did not show upregulation of KLRG1 (Fig. 1E). This documents that the preferential formation of short-lived effector T cells is specific to LCK deficiency and not a common feature of suboptimal TCR responses, such as those to suboptimal antigens.

We next monitored the presence of WT, *Lck*^-/-^, and *Lck*^CA/CA^ OT-I progeny in the blood from the peak of expansion to the memory phase. At all time points, the frequency of *Lck*^-/-^ progeny was lower than that of *Lck* WT and *Lck*^CA/CA^ counterparts (Supplemental Fig. 2D). However, the reproducibility of this observation decreased in time as memory T cells were undetectable even in some WT OT-I cells. The analysis of the progeny of the OT-I cells on day 30 post-infection showed a similar result. We observed slightly higher frequencies of KLRG1^+^ and KLRG1^+^ CD62L^+^ cells in the *Lck*^-/-^ OT-I progeny than in the WT and *Lck*^CA/CA^ OT-I progeny suggesting an effector memory phenotype (Supplemental Fig. 2E). While these results were essentially in line with the results on day six post-infection, the level of mouse-to-mouse variability was too high to make a clear conclusion concerning the effect of LCK deficiency on the phenotype of memory CD8^+^ T cells.

Overall, these data suggested that LCK-deficient T cells do respond to high-affinity cognate antigens in vivo with a reduced expansion, but with preferential expression of effector markers, especially KLRG1, in their progeny. This differentiation profile seems to be specific to the lack of LCK, since it does not mimic the outcome of suboptimal TCR signaling induced by low-affinity antigens in vivo.

### Single cell gene expression profiling reveals the effector bias of LCK-deficient T cells

To assess the role of LCK in T-cell differentiation in greater detail, we performed single-cell transcriptomics (scRNAseq) of WT, *Lck*^CA/CA^, and *Lck*^-/-^ OT-I CD8^+^ T cells responding to LM-OVA on day five post-infection. We identified five cell clusters (Fig. 2A), which were annotated as early activated, memory, effector precursors, and two clusters of effector CD8^+^ T cells (Fig. 2A-B). Whereas WT and *Lck*^CA/CA^ T cells were indistinguishable, *Lck*^-/-^ T cells were largely distinct, as they formed more effector T cells and lower frequency of memory T cells (Fig. 2C-D). Furthermore, *Lck*^-/-^ T cells displayed unique gene expression signature within the effector T-cell clusters characterized by the upregulation of effector-associated genes and specifically cytotoxic genes (*Gzmb, Prf1*), NK receptors (*Klrg1, Klrk1*), *Sell* (encoding CD62L), interferon-γ receptor (*Ifngr1*) and interferon-response genes (Fig. 2E-F, Fig. S3A-C). Moreover, *Lck*^-/-^ effector OT-I T cells downregulated the inhibitory receptors *Cd5* and *Cd6* (Fig. 2F) and upregulated genes involved in TCR signaling (Fig. 2G, Supplemental Fig. 3D), including genes encoding proximal TCR signaling components such as *Fyn, Lat, Cd8a, Cd8b*.*1*, and *Itk*, which might partially compensate for the LCK deficiency. We then used flow cytometry to verify the upregulation of GZMB as a key effector molecule in the *Lck*^-/-^ OT-I T cells at the protein level on day six post-infection (Fig. 2H).

**Figure 2.**
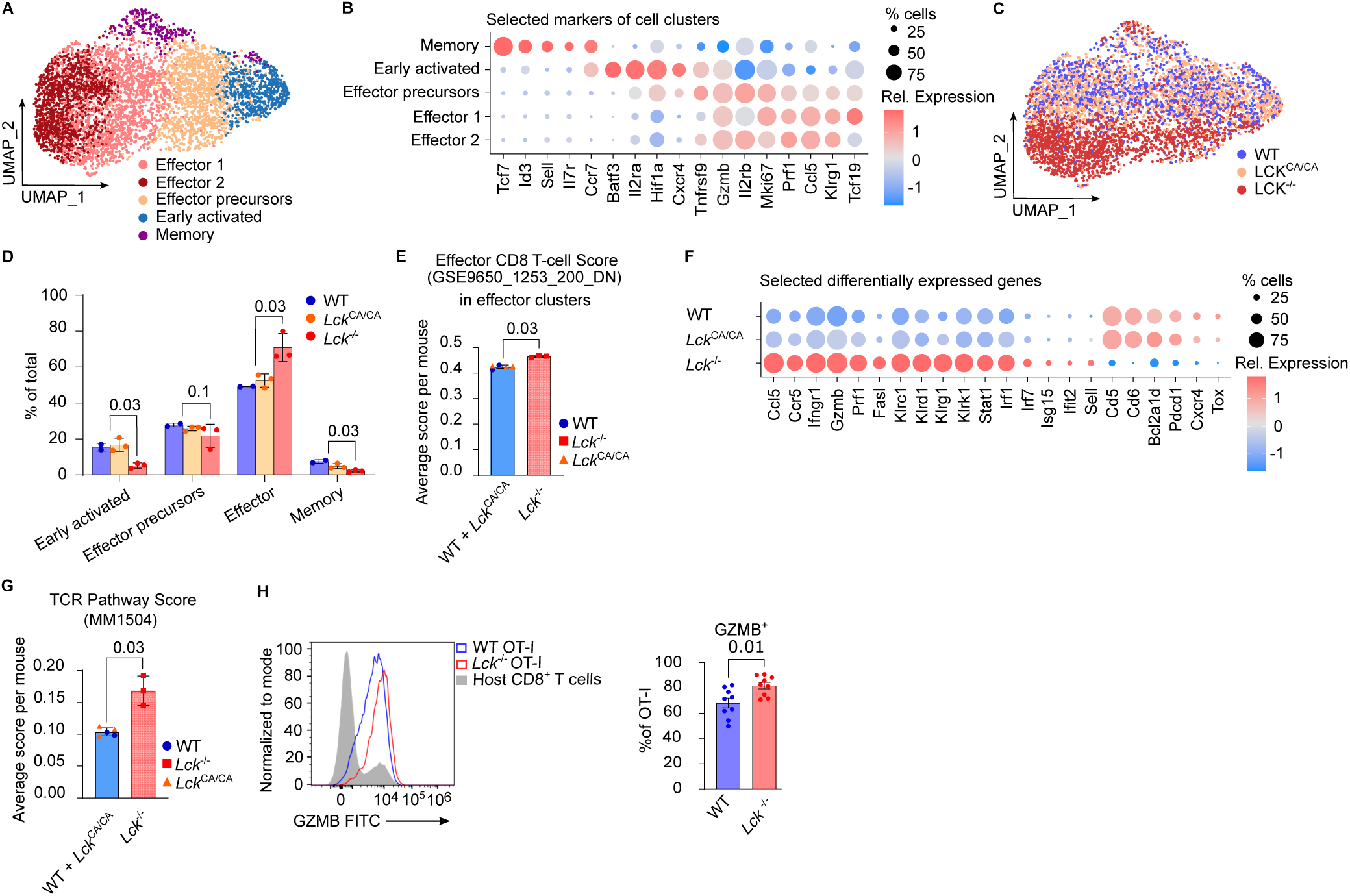
Single-cell transcriptomic analysis reveals an effector bias in *Lck*^-/-^ T cells activated in vivo. (A-G) 1×10^4^ T cells from WT, *Lck*^CA/CA^, or *Lck*^-/-^ OT-I *Rag2*^-/-^ mice were FACS-sorted as CD8α^+^ CD5^+^ and adoptively transferred to Ly5.1 host mice that were subsequently infected with LM-OVA. (A-G) OT-I CD8^+^ T cells were enriched by magnetic sorting for CD8^+^ and FACS sorted as CD8^+^ CD45.2^+^ CD45.1^-^ on day five post-infection and analyzed by single-cell RNA sequencing. n = 2 mice per group for WT, n = 3 mice for *Lck*^CA/CA^ and *Lck*^-/-^ in a single experiment. (A) UMAP projection of all OT-I T cells with annotated and color-coded clusters. The annotations were based on marker gene expression shown in (B). (B) Expression of selected marker genes used for subset annotation. Dot size and color represent the percentage of cells with detected expression of the gene and average log-normalized expression level, respectively. (C) UMAP projection of all OT-I T cells with annotated and color-coded WT, *Lck*^CA/CA^, and *Lck*^-/-^ OT-I T cells. (D) The frequency of indicated cell clusters among WT, *Lck*^CA/CA^, and *Lck*^-/-^ T cells (mean ±SEM). The two effector clusters were merged for this analysis. (E) The effector CD8^+^ T cell module score based on GSE9650 gene set (Naïve vs Eff CD8 Tcell DN gene set - M5820, human MSigDB collection) in the effector clusters of indicated OT-I genotypes. Each point represents a single mouse. WT and *Lck*^CA/CA^ mice were merged for the statistical analysis. (F) Expression of selected genes in all OT-I CD8^+^ T cells grouped by indicated genotypes. Dot size and color represent the percentage of cells with detected expression of the gene and average log-normalized expression level, respectively. (G) The Biocarta TCR Pathway gene set (MM1504, mouse MSigDB collection) module score in indicated genotypes. Each point represents a single mouse. (H) GZMB expression in WT and *Lck*^-/-^ OT-I was examined by flow cytometry on day six post-infection. A representative histogram and an aggregate analysis. n = 9 mice per group in 3 independent experiments. Mean±SEM is shown. Statistical significance was calculated using two-tailed Mann-Whitney test.

Overall, these data revealed a preferential formation of effector progeny by *Lck*^-/-^ CD8+ T cells during their response to a cognate pathogen.

### LCK-deficient effector T cells infiltrate the site of inflammation

To investigate whether LCK-deficient CD8^+^ T-cell migration to the site of infection was affected, we transferred WT, *Lck*^CA/CA^, and *Lck*^-/-^ OT-I T cells to congenic Ly5.1 mice, which were infected intranasally with influenza A virus (IAV). On day eight post-infection, *Lck*^-/-^ OT-I T cells showed a lower level of expansion and higher expression of effector markers KLRG1 and KLRK1 than *Lck* WT and *Lck*^CA/CA^ OT-I T cells in the spleen (Fig. 3A), which essentially phenocopied the results from the LM-OVA infection. In the lungs, the relative abundance of *Lck*^-/-^ OT-I T cells, measured as the percentage of all CD8^+^ T cells, was only slightly decreased in comparison to *Lck* WT OT-I T cells (Fig. 3B). The expression of KLRG1 and KLRK1 was elevated in *Lck*^-/-^ OT-I cells. Overall, these data show that LCK is dispensable for the migration of effector CD8^+^ T cells to the lungs during IAV infection.

**Figure 3.**
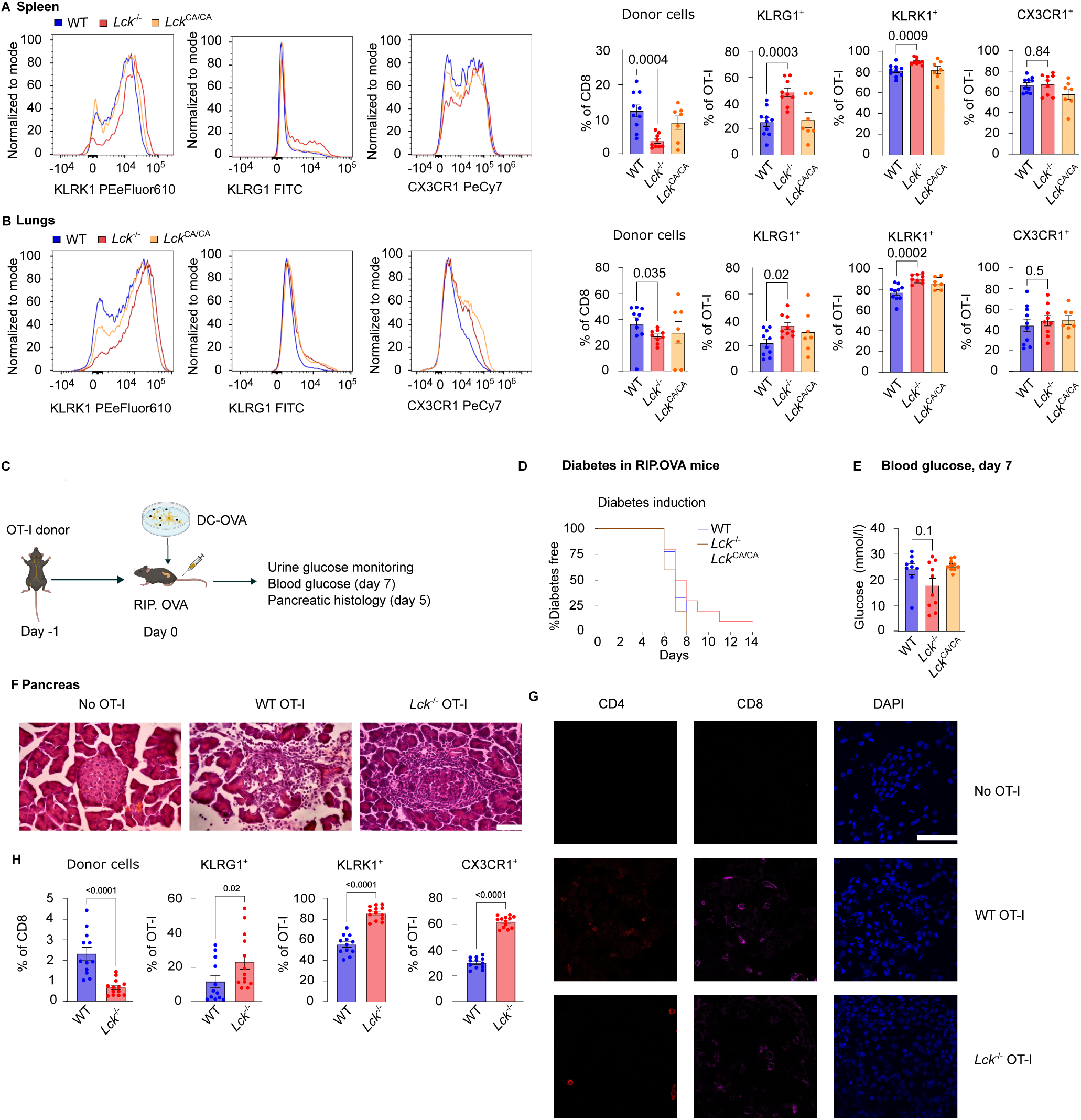
LCK-deficient CD8^+^ T cells elicit tissue-specific responses to antigens. (A-B). 1×10^4^ OT-I T cells were FACS-sorted as CD8α^+^CD5^+^ from *Lck*^WT/WT^, *Lck*^CA/CA^, or *Lck*^-/-^ OT-I *Rag2*^-/-^ mice and adoptively transferred to Ly5.1 host mice that were infected with Influenza A (Puerto Rico/8/34) expressing ovalbumin. OT-I T cells in the spleens (A) and lungs (B) were analyzed by flow cytometry on day eight post-infection. Representative histograms and aggregate results show the frequency of transferred OT-I T cells and their KLRK1, KLRG1, and CX3CR1 expression. n = 10 for WT, 9 for *Lck*^-/-^, and 8 for *Lck*^CA/CA^ mice in 3 independent experiments. (C-H). 1×10^3^ OT-I T cells were FACS-sorted as CD8α^+^CD5^+^ from the indicated mice and adoptively transferred to RIP.OVA (C-G) or Ly5.1 (H) host mice that were subsequently immunized with 1×10^6^ OVA peptide-loaded bone-marrow derived dendritic cells. (C) A schematic representation of the experiment. (D) A Kaplan-Meier curve showing the incidence of diabetes in time. The diabetes was diagnosed based on the daily measurement of glucose concentration in urine. (E) Concentration of glucose in the blood on day seven post immunization. (D-E) n = 9 for WT, 10 for *Lck*^-/-^ and *Lck*^CA/CA^ mice in 3 independent experiments. (F) Representative H&E histological analysis of pancreatic beta cells of RIP.OVA host mice stained by day six post immunization. The mice without the OT-I T cells adoptive transfer served as a negative control. The scale bar represents 50 μm. (G) Representative immunofluorescence analysis of pancreases from RIP. OVA host mice on six days post immunization. Cryosections were stained with indicated antibodies and DAPI (nuclei) and analyzed by confocal fluorescence microscopy. The scale bar represents 50 μm. (F-G) A representative image per condition out of 2 mice (no OT-I transferred control), 5 (WT), or 6 (*Lck*^*-* /-^) from two independent experiments in total. (H) Splenocytes of Ly5.1 host mice were analyzed by flow cytometry on day seven post immunization to reveal the frequency of OT-I T cells and their expression of KLRK1, KLRG1, and CX3CR1. n = 12 for WT, 13 for *Lck*^-/-^ in 3 independent experiments. Mean±SEM is shown. Statistical significance was calculated using a Mann-Whitney test.

We next compared the function of WT, *Lck*^CA/CA^, and *Lck*^-/-^ T cells in an experimental model of autoimmune type I diabetes (*21*). This model is based on the adoptive transfer of OT-I T cells to transgenic RIP.OVA mice expressing ovalbumin under the insulin promoter, followed by immunization with bone marrow-derived dendritic cells (BMDC) loaded with the OVA peptide (Fig. 3C). OT-I T cells primed by these BMDCs proliferate, form cytotoxic effector T cells, infiltrate the pancreas and kill insulin-producing β cells. Both the initial activation and the target cell killing require TCR signaling. We previously titrated the numbers of transferred OT-I T cells, revealing that 1,000 WT OT-I cells were sufficient to induce diabetes in most mice, whereas 250 OT-I cells were tolerated in the majority of mice (*22*). Here, the transfer of 1,000 WT or *Lck*^CA/CA^ OT-I T cells induced diabetes in all mice by day eight. The same number of *Lck*^-/-^ OT-I cells induced diabetes in 90% of the mice albeit with a slight delay compared to WT OT-I cells (Fig. 3D-E). The histological analysis of the pancreas revealed the destruction of the pancreatic islets and immune infiltration, including CD4^+^ and CD8^+^ T cells, in mice with a previous transfer of WT or *Lck*^-/-^ OT-I cells, but not in mice without any OT-I transfer (Fig. 3F-G). The in vivo response of OT-I T cells to BMDC-OVA priming induced a relatively weaker expansion of *Lck*^-/-^ OT-I T cells, but a preferential differentiation into effector T cells (Fig. 3H, Supplemental Fig. 4A), which explains their ability to induce the autoimmune diabetes.

Taken together, the IAV and T1D models demonstrated that preferential differentiation of *Lck*^-/-^ CD8^+^ T cells into tissue-infiltrating effector T cells partially compensated for their reduced proliferative response.

### Downstream TCR signaling pathways are differentially affected by LCK deficiency

In the next step, we analyzed which downstream signaling pathways are most affected by LCK deficiency using ex vivo activation assays. To detect signaling intermediates by flow cytometry, we used T2-Kb cells loaded with OVA peptide as APCs (Fig. 4A). We observed a strong reduction in the phosphorylation of ERK1/2, a mild reduction in IκB degradation, and only minimally affected cJUN and S6 phosphorylation in the *Lck*^-/-^ OT-I T cells. These data suggest that particular downstream signaling pathways have differential sensitivity to LCK deficiency (Fig. 4B-C). We then analyzed the nuclear translocation of NFAT and NFκB following OVA peptide stimulation by imaging cytometry (Fig. 4D-E, Supplemental Figure 4B). We observed that the translocation of both transcription factors was reduced in the *Lck*^-/-^ OT-I T cells, although the translocation of NFκB was hardly detectable even in the WT cells in this assay. Overall, LCK deficiency on downstream TCR signaling selectively impacts the ERK and the NFAT pathways.

**Figure 4.**
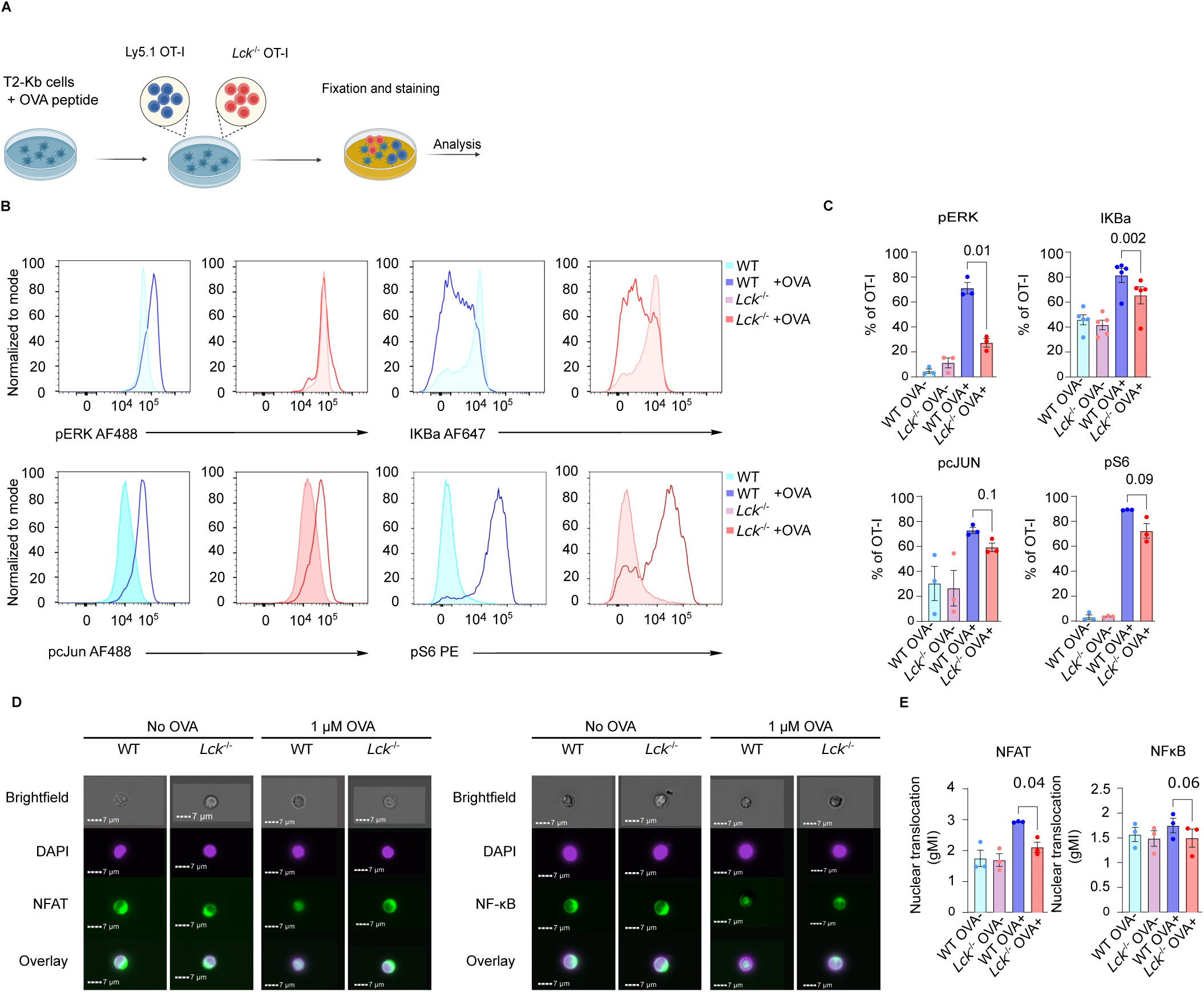
LCK differentially regulates specific downstream TCR signaling pathways. (A-E) Cells isolated from spleens and lymph nodes of Ly5.1 WT OT-I *Rag2*^-/-^ and Ly5.2 *Lck*^-/-^ OT-I *Rag2*^-/-^ mice were mixed at 1:9 (B-C) or 2:8 (D-E) ratio and co-cultured with OVA peptide-loaded T2-Kb cells at 2:1 ratio (B-C) or activated with 1 µM OVA peptide (D-E) for 1 hour. (A) A schematic illustration of the experiments with T2-Kb cells. (B-C) Signaling intermediates were quantified by flow cytometry. Representative histograms (B) and aggregate results of the percentage of positive cells (C) are shown. n = 5 independent experiments for IKBa or 3 for other signaling intermediates. (D-E) OT-I T cells were activated or not and stained for CD8, TCRβ, DAPI and NFAT or NFκB. Nuclear translocations of NFAT and NFκB were analyzed by imaging flow cytometry (for gating strategy see Fig. S4C). Representative images (D) and the aggregate data of the nuclear translocation scores (E) are shown. Scale bar 7 µm. n = 3 independent experiments. Mean±SEM. Statistical significance was determined using two-tailed paired t test.

### Deletion of LCK in mature OT-I T cells

The above-described experiments suggested that *Lck*^-/-^ CD8^+^ T cells exhibit relatively strong antigenic responses in vivo, which contradicts the common view of LCK as the essential proximal TCR kinase. To further verify our previous results, we employed another experimental model based on an acute deletion of *Lck* in mature non-activated OT-I cells using CRISPR-Cas9 nucleofection (*23*) (Fig. 5A). OT-I cells receiving three *Lck-*targeting crRNAs were designated Lck KO, to distinguish them from T cells isolated from the germline *Lck*^*−/−*^ mice described above. Since this type of gene targeting is not efficient in all cells, we expected the *Lck* KO cells to constitute a mixture of cells with and without *Lck* disruption. However, the percentage of cells with the *Lck* disruption could not be determined immediately after the nucleofection by flow cytometry, as the pre-existing LCK protein would still be present in these cells. Control cells received a non-targeting crRNA. After the nucleofection, 1×10^5^ the *Lck* KO and control OT-I cells were adoptively transferred into congenic Ly5.1 host mice that were infected with LM-OVA on the next day. The rest of the cells were activated in vitro in an LCK-independent manner by PMA/ionomycin for three days and then probed for intracellular LCK presence by flow cytometry to estimate the knock-out efficiency.

**Figure 5.**
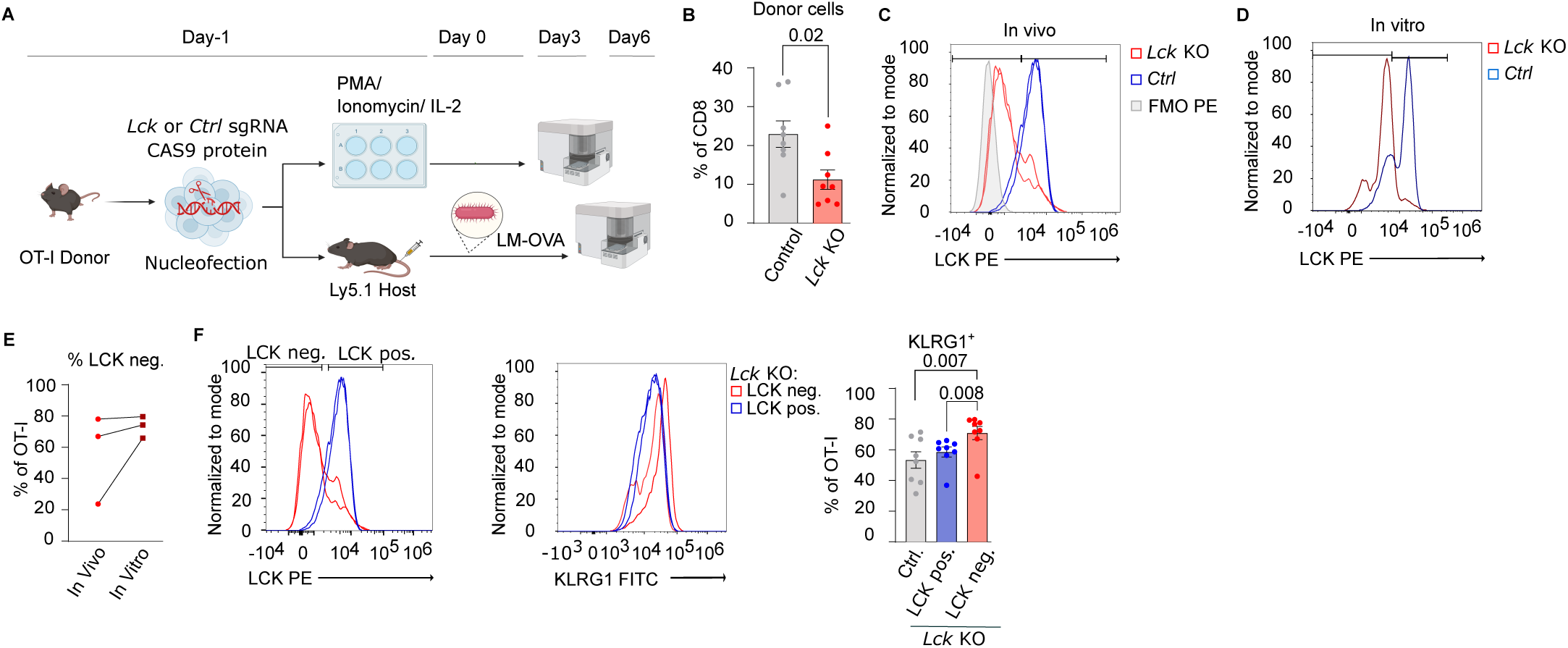
LCK knock-out in mature CD8^+^ T cells leads to KLRG1 upregulation upon in vivo activation. (A-F) WT OT-I T cells were nucleofected with recombinant CAS9 and *Lck*-targeting (*Lck* KO) or mock (Ctrl.) crRNAs. Then the cells were split. 1×10^5^ cells per mouse were adoptively transferred to Ly5.1 host mice subsequently infected with LM-OVA and analyzed by flow cytometry on day six post-infection (B, D-E, F). 1.5×10^6^ OT-I T cells activated with PMA/Ionomycin for 24 h and then cultured in IL-2-supplemented medium for additional 48 h and analyzed by flow cytometry (B-F). n = 8 mice per group in 3 independent experiments. (A) A schematic illustration of the experiment. (B) Bar graph showing the percentage of adoptively transferred *Lck* KO and Ctrl. OT-I cells out of CD8^+^ six days after LM-OVA infection. Mean±SEM. Statistical significance was calculated using two-tailed Mann-Whitney test. (C-D) Representative histograms showing the efficiency of the *Lck* KO in the in vivo responding cells (C) and ex vivo cultured cells (D). (E) The comparison of LCK^-^ T cells in the *Lck* KO cells *in vivo* and *in vitro*. Mean of 2-3 mice per group were used for the *in vivo* part of the experiment. n = 3 independent experiments. (F) Representative histograms and aggregate results showing the gated LCK negative (neg.) and LCK positive (pos.) cells in vivo and the expression of KLRG1 on LCK pos. and LCK neg in *Lck* KO and control T cells. Mean±SEM. Statistical significance was calculated using two-tailed paired Wilcoxon signed-rank test (LCK neg. vs. LCK pos.) or two tailed Mann-Whitney test (LCK neg. vs Ctrl).

We observed the *Lck* KO OT-I cells expanded less than control OT-I cells on day six post-infection, but the difference was relatively subtle. Using flow cytometry, we detected a mixture of LCK^+^ and LCK^-^ (successful KO) donor OT-I cells in hosts which have received the *Lck* KO cell mixture on day six post-infection. The fact that we detected LCK^-^ OT-I cells indicated that these cells expanded during their response to their cognate antigen in this experimental setup (Fig. 5C). The in vitro PMA/ionomycin-activated T cells, which expanded in a TCR-independent way, showed only slightly higher percentage of LCK^-^ cells among the *Lck* KO (Fig. 5D) indicating that the proliferative disadvantage of *Lck* KO OT-I in vivo was relatively minor (Fig. 5 C,E). KLRG1 expression on OT-I T cells was analyzed using stricter gating of LCK^+^ and LCK− populations to minimize spillover between gates and avoid potential heterozygous cells with only partially reduced LCK expression (Fig. 5F). The LCK^-^ OT-I cells responding to the LM-OVA infection in vivo showed higher expression of KLRG1 in comparison to LCK^+^ positive OT-I cells in the same host as well as in comparison to control OT-I cells (Fig. 5F), indicating preferential effector T-cell formation in the absence of LCK. Overall, these data showed that key features observed in OT-I T cells isolated from *Lck*^-/-^ mice with a germ-line *Lck* deletion, i.e., the ability of LCK-deficient T cells to respond to a cognate antigen in vivo and their enhanced effector T-cell formation, are reproduced in the experimental setup with acute LCK deletion in mature T cells.

### Analysis of FYN-deficient T cells

Previous data suggested that FYN partially takes over the function of LCK in *Lck*^-/-^ thymocytes and T cells (*4, 5, 15*). Accordingly, we observed a slight upregulation of FYN in *Lck*^-/-^ compared to *Lck* WT cells after activation and at the steady state (Supplemental Fig. 5A-C). For this reason, we generated *Fyn*^-/-^ mice (Fig. 6A). First, we analyzed the recruitment of FYN to the TCR/CD3 complex using a newly designed proximity ligation assay between FYN and CD3ε. Using *Fyn*^-/-^ T cells as a baseline control, we investigated the proximity of FYN and the TCR/CD3 (Supplemental Fig. 5D). The data did not show significantly enhanced FYN-TCR/CD3 proximity in *Lck*^-/-^ T cells, neither in unstimulated condition nor after anti-CD3 activation (Supplemental Fig. 5D). Altogether, these experiments show that Fyn is not strongly recruited to the TCR upon activation in contrast to the behavior reported for LCK (*24, 25*).

**Figure 6.**
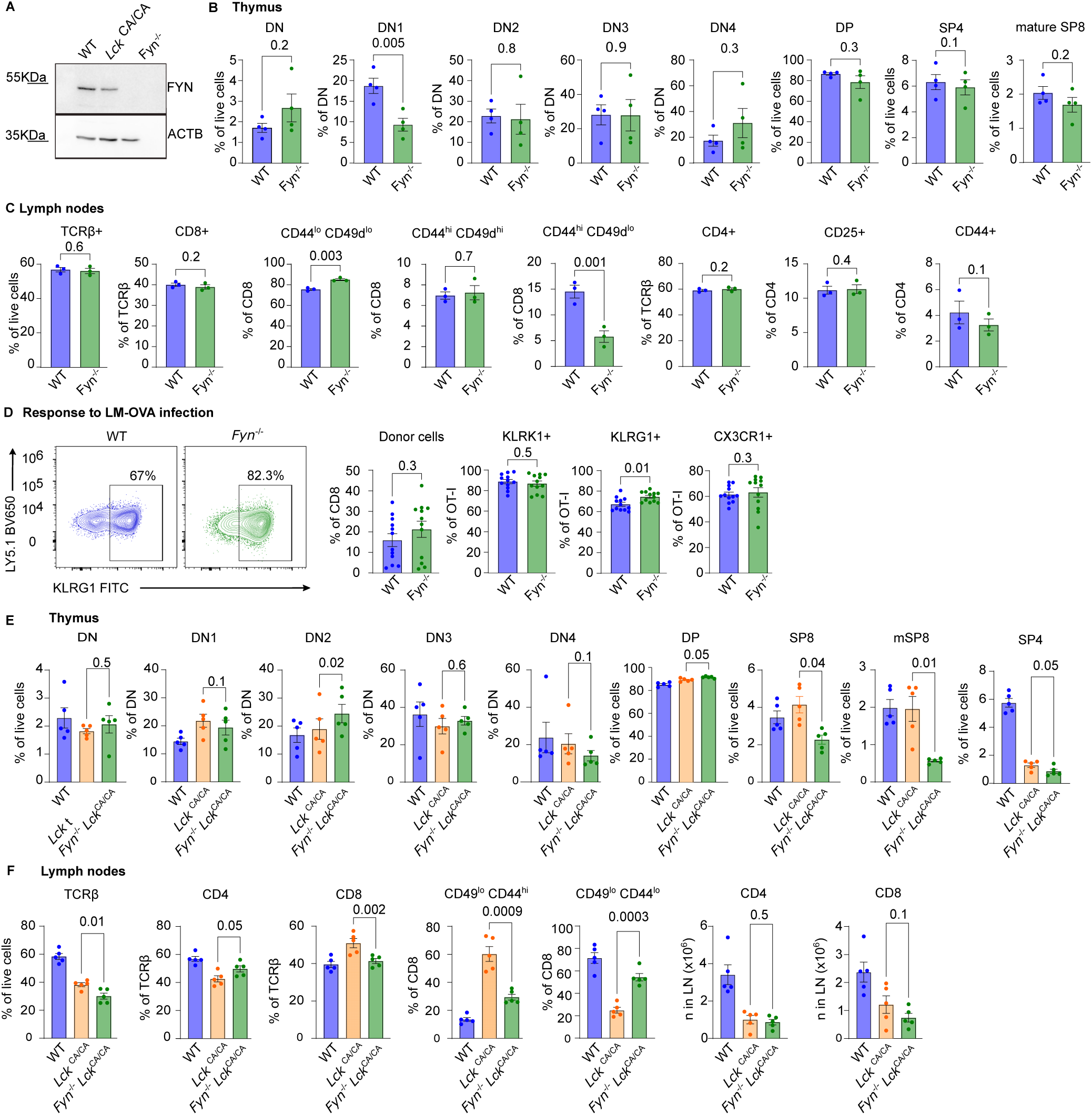
Phenotype of *Fyn*^-/-^ deficient T-cells mice at the steady state and during the immune response. (A) WT, *Lck*^CA/CA^, and *Fyn*^-/-^ splenocyte lysates were immunoblotted using antibodies against FYN (59 KDa) and ACTB (40 KDa, loading control). n = 2 independent experiments. (B-C) The aggregate results of indicated T-cell subsets in the thymus (B) and lymph nodes (C) of WT and *Fyn*^-/-^ mice. n = 4 mice in 4 independent experiments (B) or n = 3 mice in 3 independent experiments (C). Mean±SEM. Statistical significance was calculated using two-tailed paired t test (D) 1×10^4^ T cells from WT OT-I *Rag2*^-/-^ or *Fyn*^-/-^ OT-I *Rag2*^-/-^ mice were adoptively transferred to Ly5.1 host mice that were subsequently infected with LM-OVA. Splenocytes were analyzed by flow cytometry on day six post-infection. Representative contour plots and aggregate results show the frequency of OT-I T cells and the expression of indicated markers. n = 12 mice per group in 3 independent experiments. Statistical significance was calculated using two-tailed Mann-Whitney test. (E) The aggregate results of indicated T-cell subsets in the thymus (B) and lymph nodes (C) of WT, *Lck*^CA/CA^, *and Lck*^CA/CA^ *Fyn*^-/-^ mice. n = 5 mice in 5 independent experiments. Mean±SEM. Statistical significance was calculated using two-tailed paired t test.

We hypothesized that the preferential effector phenotype of *Lck*^-/-^ OT-I might be caused either by the overall lower SFK activity (LCK and FYN combined) or by intrinsic non-overlapping roles of LCK and FYN. T-cell development was normal both in polyclonal and monoclonal *Fyn*^-/-^ OT-I mice (Fig. 6B-C, Supplemental Fig. 5E-F). In response to LM-OVA infection, *Fyn*^-/-^ OT-I cells expanded to a similar extent as WT OT-I T cells, yet exhibited a modestly higher frequency of KLRG1^+^ effector T cells. (Fig. 6D). These data suggested that the reduction of the combined activity of LCK and FYN in *Lck*^-/-^ and, to a lesser extent, in *Fyn*^-/-^ T cells stimulate an increased formation of short-lived effector KLRG1^+^ T cells, revealing a regulatory role of these kinases in T-cell activation.

Finally, to investigate whether FYN is dispensable for T-cell development in the condition of disrupted LCK-co-receptor interaction, we analyzed T-cell development in *Fyn*^-/-^ *Lck*^CA/CA^ mice. Whereas *Lck*^CA/CA^ mice show a severe impairment of CD4^+^ T-cell formation, the generation of CD8^+^ T cells was much less affected (*3*) (Fig. 6E-F). However, the *Fyn*^-/-^ *Lck*^CA/CA^ double transgenic mice showed strongly impaired formation of both CD4^+^ and CD8^+^ T cells indicating that the relative independence of the CD8^+^ lineage of the CD8-LCK interaction requires the activity of FYN. These data show an additional previously not appreciated level of LCK and FYN interplay, which is unmasked in the mice with disrupted CD8-LCK interaction.

## Discussion

In this study, we show that LCK is dispensable for primary responses of murine CD8^+^ T cells in vivo. Our findings contrast with observations in human Jurkat T-ALL cells (*3, 26*), where LCK deficiency completely abolishes downstream TCR signaling, and in mouse thymocyte development, where LCK deficiency leads to a severe block in T-cell maturation (*3, 6*). However, our data align with previous results indicating that *LCK*^-/-^ human T cells expressing CARs are actually more efficient than their WT counterparts (*14*) and with findings that LCK is dispensable for secondary responses of memory CD8^+^ T-cells in mice (*13*). These two studies suggested that LCK is not essential for antigenic responses in mature T cells. Our work uncovers, for the first time, a specific role for LCK for TCR signaling depending on the T-cell maturation stage. This finding also helps to explain why patients with genetic *LCK* deficiency have at least partially preserved cytotoxic functions of CD8^+^ T cells in the periphery. Still, the key role of LCK in thymic T-cell maturation has a negative impact on the peripheral CD8^+^ T cell numbers and biases the repertoire of these cells towards higher self-reactivity. This differential role of LCK in thymic and peripheral T cells might contribute to the autoimmune symptoms reported in some LCK-deficient patients (*9*).

Whereas weak TCR signaling induced by low-affinity antigens leads both to weak proliferation and low level of effector cell formation in CD8^+^ T cells (*20*), LCK deficiency selectively impacts CD8^+^ T cell proliferation. Antigen-induced proliferation was reduced in *Lck*^-/-^ CD8^+^ T cells in vivo, consistent with prior in vitro findings using allogeneic stimulation (*16*) or anti-CD3 antibody (*15*). In contrast, we observed that *Lck*^-/-^ CD8^+^ T cells have a skewed differentiation program forming cytotoxic effector T-cell progeny at even slightly higher frequency in comparison to WT T cells. Accordingly, *Lck*^-/-^ CD8^+^ T cells induced experimental autoimmune diabetes in a mouse model, demonstrating their capacity to complete all steps of the effector program, including priming, expansion, tissue infiltration, and β-cell destruction. These results are in line with a previous study showing that murine LCK-deficient T cells show normal killing of allogenic targets in vitro (*16*), but contrast with another study showing a key role of LCK in target cell killing in an antigen-dependent manner in vitro (*17*). These conflicting data indicate that the effector function of T cells in vitro strongly depends on protocol used and that in vivo experiments are crucial for evaluating the role of particular signaling proteins for T-cell effector functions.

A previous study showed that primary CD8^+^ T cell anti-viral responses are fully LCK-dependent, since they did not identify any tetramer-specific CD8^+^ T cells in vaccinia or LCMV-infected LCK-deficient mice over the background (*13*), which disabled a subsequent analysis of their memory or effector phenotype. Tewari et al. used *Lck*^-/-^ mice expressing a doxycycline-driven LCK transgene, where the LCK deficiency was induced by doxycycline withdrawal (*13*). In these mice, all CD8^+^ and CD4^+^ T cells lacked LCK. In contrast, we studied intrinsic role of LCK in CD8^+^ T-cells with fixed OT-I TCR adoptively transferred to polyclonal WT mice with endogenous LCK-expressing T cells using two complementary approaches: whole body *Lck*^-/-^ mice or *Lck* KO in mature T cells. Our model systems investigated the intrinsic role of LCK in CD8^+^ T-cell responses on a per-cell basis, as it preserved the competition with LCK-expressing T cells on the one hand and the by-stander help of host CD4^+^ and CD8^+^ T cells (e.g., via IL-2 production) on the other hand. For this reason, our study is especially relevant for adoptive T-cell therapies where LCK levels can be modulated, such as the use of LCK-deficient T cells expressing CARs (*14*).

Our signaling analyses indicate that triggering of the TCR in *Lck*^-/-^ deficient T cells was probably mediated by FYN, as indicated by slight upregulation of FYN in these cells and previous reports showing the ability of FYN to partially compensate LCK deficiency (*4, 5, 16*). *Fyn*^-/-^ T cells showed normal T-cell development and peripheral antigenic responses with a slight bias towards the formation of KLRG1^+^ effector progeny. These results suggested that the reduction of the overall pool of SFK activity enhances effector formation, probably via dysregulation of downstream activation pathways. Reduced activation of inhibitory receptors such as PD-1 and CTLA4, along with lower levels of inhibitory receptors CD5 and CD6 in *Lck*^-/-^ T cells may further contribute to enhanced effector differentiation (*27-30*).

The analysis of downstream TCR signaling pathways in *Lck*^-/-^ T cells triggered by the cognate antigens revealed a strong reduction in the calcium-dependent NFAT nuclear translocation and ERK1/2 activation by phosphorylation. Accordingly, the defect in calcium influx has been described upon stimulation of LCK-deficient T cells using anti-CD3 (*16*) or anti-CD3 and anti-CD4 (*15*) antibodies and impaired ERK1/2 phosphorylation was observed in antigen-stimulated F5 TCR transgenic CD8^+^ T cells in vitro (*15*). In contrast, the NFκB, AKT/mTOR and JNK/AP-1 pathways were less affected by LCK deficiency. Overall, LCK deficiency leads to rewiring of downstream signaling pathways in CD8^+^ T cells, which causes intrinsically reduced proliferation and enhanced cytotoxic effector formation on a per cell basis in vivo.

Altogether, our and previous data indicate that FYN partially takes over the role of LCK in *Lck*^-/-^ T cells depending on the T cell maturation stage. We have previously generated a knock-in mice in which LCK interaction with CD4 and CD8 co-receptors was disrupted (*3*). This mouse revealed an impaired formation of mature CD4^+^ thymocytes and peripheral T cells, whereas the formation of CD8^+^ T cells was less affected. Here, we show that the development of CD8^+^ T cells in these mice largely depends on FYN activity, as *Lck*^CA/CA^ *Fyn*^-/-^ mice show a strong block in the formation of mature CD8^+^ single positive thymocytes and peripheral T cells. It is unclear whether the overall SFK activity towards the TCR complex drops below a critical threshold in these double transgenic mice or whether there is a specific interplay between these two kinases in thymocytes, as was proposed by some older studies (*31, 32*). Previous studies showed that *Fyn*^-/-^ thymocytes and T cells have impaired responses to TCR cross-linking via anti-CD3 antibodies (*11, 33*), whereas their responses to cognate antigens, staphylococcal enterotoxin A super-antigen, and allogeneic stimulation are intact (*11, 33*). This suggests that FYN contributes to TCR signaling when co-receptors are not involved, a situation that likely mirrors the absence of co-receptor– bound LCK in the *Lck*^CA/CA^ mice. Moreover, thymocytes were shown to be more FYN-dependent than mature T cells in terms of TCR signaling (*33*), which might relate to the defect in thymocyte maturation in the *Lck*^CA/CA^ *Fyn*^-/-^ mice. Taken together, FYN plays a key role in the development of CD8^+^ T cells when LCK lacks its ability to interact with co-receptors.

## Methods

### Mice

Experimental mice were 6 to 12 weeks old at the beginning of the experiments, except for steady-state phenotyping studies, which employed mice aged 4 to 6 weeks. Both male and females were included in all experiments, with littermates used when possible. We aimed to have representative population of both sexes among experimental groups in all experiments.

Mice were maintained in specific pathogen-free facility (Institute of Molecular Genetics of the Czech Academy of Sciences; IMG). Mice were kept in the facility with a 12-h light/12-h dark cycle and temperature and relative humidity maintained at 22⍰±⍰1⍰°C and 55⍰±⍰5%, respectively. Mice were fed with standard rodent breeding diet and quenched with filtered water ad libitum. Animal protocols used were in accordance with the laws of the Czech Republic and approved by the Resort Professional Commission for Approval of Projects of Experiments on Animals of the Czech Academy of Sciences, Czech Republic (ID AVCR 72/2018, AVCR 2378/2022 SOVII, AVCR 1667 2022 SOV II, and AVCR 5262/2024 SOV II).

The following mice had C57BL/6J background: Ly5.1 (RRID: IMSR_JAX:002014) (*34*), OT-I *Rag2*^-/-^ ((RRID:MGI:3783776, MGI:217491) (*35, 36*) RIP.OVA (RRID:IMSR_JAX:005433) (*37*), *Lck*^C20.23A/C20.23A^ (*3*), *Lck*^-/-^ (*38*).

*Fyn*^-/-^ (MGI: Fyn^em1(IMPC)Ccpcz^) mice were generated on C57BL/6N background by targeting exon 4 (ENSMUSE00000407193) in the Czech Centre for Phenogenomics, Institute of Molecular Genetics, CAS. A frame-shift deletion was generated using CRISPR/Cas9 technology. The guide RNAs (gRNAs) with the highest score and specificity were designed using http://crispor.tefor.net/. The following guides were selected: gRNA1-GATGGAGGTCACGCCGAAGC and gRNA2-CAGGGACTCACCGTCTTTGG. The gRNAs (80 ng/µl) were assembled into a ribonucleoprotein (RNP) complex with Cas9 protein (200 ng/µl; 1081058, 1072532, Integrated DNA technologies), electroporated into 1-cell zygotes, and transferred into pseudopregnant foster mice. Putative founders were analyzed by PCR and sequencing. A founder harboring a 68 bp deletion, affecting part of exon 4, was chosen for subsequent breeding. Genotyping was performed by PCR with forward (F) 5⍰-TTGGCCGAGGACTGAGATAC-3⍰ and reverse (R)5’-TCTCCGATTACCACACACCTG-3⍰ primers. The F and R products are 507 bp in the wild-type and 439 bp in the mutant animal. A selected founder was bred with C57BL/6NC wild-type to confirm germ-line transmission of the target deletion. The established strain is available in the EMMA/Infrafrontier repository CCP node.

*Fyn*^-/-^ were further (back)crossed to OT-I *Rag2*^-/-^ strain(s), which were on the C57BL/6J background. Littermates were used for analysis.

### Cell culture and counting

Cells were counted by using Cytek Aurora flow cytometer (Cytek), Z2 Coulter Counter Analyzer (Beckman Coulter), or LUNA-II™ Automated Cell Counter.

Dendritic cells were generated from the bone marrow extracted from the tibia and femur of female Ly5.1 or C57BL/6J mouse between 6 to 8 weeks old. Stem cells derived from bone marrow were cultured on 100 mm untreated plates for 10 days. Cells were maintained in IMDM supplemented with 10% FBS (GIBCO), 100 U/ml penicillin (BB Pharma), 100 mg/ml streptomycin (Sigma-Aldrich), 40 mg/ml gentamicin (Sandoz), and 2% of supernatant from J558 cells producing GM-CSF at 5% CO2 at 37°C (*39*). The cells were split on day 3, 5 and 8. On day 10, cells were activated with 100 ng/ml LPS (Sigma-Aldrich) and 200 nM of OVA peptide (SIINFEKL) for 3 h at 5% CO2 at 37 °C. Cells were incubated for 5 min at 5% CO2 at 37 °C with 0.02% EDTA in PBS and harvested. Cells were washed, filtered, counted and injected.

T2-Kb (*40*) (provided by E. Palmer, University Hospital Basel) were maintained in RPMI media (Sigma) complemented with 10% FBS/100 U/ml penicillin/100 µg/ml streptomycin/40 µg/ml gentamicin. T2-Kb cells were used as antigen presenting cells to OT-I T cells.

All the cell lines were regularly tested for mycoplasma by PCR.

### Antibodies

The following antibodies from BioLegend were used: CD4 AF647 (Cat# 100530, IgG2a, κ, Clone RM4-5), CD8a AF594 (Cat# 100758, IgG2a, κ, Clone 53-6.7), CD8a BV421 (Cat# 100753, IgG2a, κ, Clone 53-6.7), in PE (Cat# 100708, IgG2a, κ, Clone 53-6.7), CD127 BV421 (Cat# 135027, IgG2a, κ, Clone A7R34), CD62L AF700 (Cat# 104426, IgG2a, κ, Clone MEL-14), CD62L APC (Cat# 104412, IgG2a, κ, Clone MEL-14), CD44 BV785 (Cat# 103059, IgG2b, κ, Clone IM7), PerCP-Cy5.5 (Cat# 103032, IgG2b, κ, Clone IM7), BV650 (Cat# 103049, IgG2b, κ, Clone IM7), and PE (Cat# 103008, IgG2b, κ, Clone IM7), KLRG1 in FITC (Cat# 138410, IgG Clone 2F1/KLRG1), and APC (Cat# 138412, IgG, Clone 2F1/KLRG1), CD25 BV785 (Cat# 102051, IgG1, λ, Clone PC61), BV650 (Cat# 102038, IgG1, λ, Clone PC61), BV605 (Cat# 102036, IgG1, λ clone PC61), CD25 PE-Cy7 (Cat# 102016, IgG1, λ, Clone PC61), CD49d Pe-Dazzle594 (Cat# 103625, IgG2b, κ, Clone R1-2) and APC (Cat# 103622, IgG2b, κ, Clone R1-2), TCRβ FITC (Cat# 109206, IgG, Clone H57-597), BV711 (Cat# 109243, IgG2, λ1, Clone H57-597), Pacific Blue (Cat# 109226, IgG, Clone H57-597), PE (Cat# 109208, IgG, Clone H57-597), AF488 (Cat# 109215, IgG, Clone H57-597), in PeCy7 (Cat# 109222, IgG, Clone H57-597), Ly5.1 in BV650 (Cat# 110735, IgG2a, κ, Clone A20), PerCP-Cy5.5 (Cat# 110728, IgG2a, κ, Clone A20), in FITC (Cat# 110706, IgG2a, κ, Clone A20), and AF700 (Cat# 110724, IgG2b, κ, CloneA20), Ly5.2 AF700 (Cat# 109822, IgG2a, κ, Clone 104), and APC (Cat# 109814, IgG2a, κ, Clone 104), PeCy7 (Cat# 109830, IgG2a, κ, Clone 104), APCCy7 (Cat# 109824, IgG2a, κ, Clone 104), CD45 FITC (Cat# 103108, IgG2b, κ, Clone 30-F11), CD45R/B220 BV510 (Cat# 103248, IgG2a, κ, Clone RA3-6B2), CD4 BV510 (Cat# 100559, IgG2a, κ, Clone RM4-5), AF700 (Cat# 100536, IgG2a, κ, Clone RM4-5), and PE (Cat# 130310, IgG2a, κ, Clone H129.19), CX3CR1 PE-Cy7 (Cat# 149016, IgG2a, κ, Clone SA011F11), CD5 APC (Cat# 100625, IgG2b, κ, Clone 53-7-3H), CD69 PE (Cat# 104508, IgG1, λ3, Clone H1.2F3), CD24 FITC (Cat# 101806, IgG2b, κ, Clone M1/69), anti-CD80 in PerCPCy5.5 (Cat#104722, isotype IgG, Clone 16-10A1), Granzyme B in FITC (Cat# 515403, IgG1, κ, Clone GB11).

The following antibodies from BD Pharmingen were used: CD4 PE (Cat# 553652, IgG2a, κ, Clone H129.19), CD5 PerCP (Cat# 553025, IgG2a, κ, Clone 53-7-3), CD8 PerCP (Cat# 553036, IgG2a, κ, Clone 53-6.7), CD49d PE (Cat# 553157, IgG2b, κ, Clone R1-2), LCK PE (Cat# 558496, IgG1, κ, clone MOL 171, 1:50), CD16/CD32 purified (Cat# 553141, isotype IgG2b, κ, 1:100), CD11c in Pe-Cy7 (Cat# 558079, isotype IgG1, λ2, Clone HL3).

The following antibodies were purchased from eBioscience: CD314 PE-eFluor610 (Cat# 61-5882-82, Clone CX5), CD119 PE (Cat# 12-1191-82,, Clone 2E2), and CD4 AF700 (Cat# 56-0042-82, IgG2a, κ, Clone RM4-5), anti-MHCII (IA/IE) in FITC (Cat# 11-5321-82, IgG2b, κ, Clone M5/114.15.2), CD86 in AF700 (Cat#105024, isotype IgG2a, κ, Clone GL-1), Perforin in PE (Cat# 12 - 9392-80, IgG2a, κ, Clone eBioOMAK-D).

The following antibodies were purchased from Cell signaling: Phospho-p44/42 MAPK XP (Cat# 4370, IgG), Phospho-c-Jun (Cat# 3270, IgG), NFAT1 XP (Cat# 5861S, IgG), IκBα in AF 647 (Cat# 8993, IgG1), Phospho-S6 Ribosomal Protein (Ser235/236) (Cat# 4858, IgG) or Santa Cruz: NFκB p65 (Cat# sc-372, clone C-20, IgG).

From Abcam CD8α primary antibody (Cat# ab217344, IgG, Clone EPR21769) was purchased.

Following primary antibodies were used: FYN (FYN-01, Invitrogen, Cat# MA119331, IgG2b, κ, 1:500 diluted), FYN (Cell signaling, Cat# 4023), anti-β-Actin (Sigma, Cat# A1978, clone AC-15), anti-CD311 (Everest #EB12592, diluted 1:400), Insulin polyclonal antibody (Invitrogen, Cat# PA1-26938, IgG, 1:50).

The following secondary antibodies were used: Goat anti-rabbit purchased from Invitrogen in AF488 (Cat# A-11008, IgG H+L, 1:1000) and in AF555 (Cat# A-32732, IgG H+L, 1:1000), anti-mouse (Mouse TrueBlot ULTRA HRP conjugated, Cat# 18-8817-33, Rockland, IgG), anti-guinea pig in AF488 (Invitrogen, Cat# A-11073, IgG H+L, 1:1000), Duolink *In Situ* PLA Probe anti-rabbit PLUS (Cat# DUO92002, Sigma, diluted 1:7), and Duolink *In Situ* PLA Probe anti-goat MINUS (Cat# DUO92006, Sigma, 1:7).

For immunofluorescence staining, pancreases were snap-frozen, cryosectioned, and subsequently stained with indicated antibodies in PBS/0.3% Triton X-100. For flow cytometry analysis of mouse blood, splenocytes, and lung samples, single-cell suspensions were stained in FACS buffer, 1:200 diluted if not written differently.

Viability dye LIVE/DEAD fixable near-IR dye (Thermo Fisher Scientific, L34975) was used for the viability staining (1:500). Biotinylated H2-K^b^-OVA monomers were obtained from the NIH Tetramer Core Facility (USA) and combined with Streptavidin, R-Phycoerythrin Conjugate (SAPE) (Invitrogen, Cat# S866) to assemble K^b^-OVA tetramers.

DAPI (ThermoFisher #D1306 or SouthernBiotech #0100-20) were used for the identification of the nuclei.

### Flow cytometry and imaging cytometry

Lymphocytes were isolated from the previously described mouse strains. Single cell suspensions were created by dissociating tissues on ice in PBS or FACS buffer (PBS, 2% FBS, +/-0.1% sodium azide) followed by filtration through 70 nm pore filter. Lungs were processed on day 8 post-influenza infection using lung dissociation kit (130-095-927, Miltenyi Biotec), according to manufactures instructions. The erythrocytes were lysed in ACK buffer for 2 min at RT. The staining of live cells with indicated antibodies and live/dead dye was performed in FACS buffer on ice for ∼30⍰min.

For the staining of Granzyme B and Perforin, BD Cytofix/Cytoperm Fixation/Permeabilization Kit was used (BD Bioscience #554714) according to manufacturer’s instructions. The cells were stained for 45 min-1 h at room temperature (RT).

For the analysis of signaling intermediates, the cells were fixed with 2-4% formaldehyde (Sigma #F8775) at RT, followed by permeabilization with 0.3% Triton X-100. Samples were stained overnight at RT with specific primary antibodies. The next day, samples were stained with a fluorescently labeled secondary antibody for 45 min.

The samples were measured on an Aurora (Cytek) or FACS Symphony (BD Bioscience) flow cytometer. The data were analyzed using Flow Jo (version 10.6.2, BD Biosciences). Cell sorting was performed using an Aria or Influx (both BD Biosciences) using a 100 μm nozzle.

The imaging cytometry was performed using an Amnis Imagestream X Mk II Imaging Flow Cytometer (Luminex). Data were analyzed using Ideas 6.2 software (Amnis). The nuclear translocation index was calculated as a Similarity feature between the channels corresponding to the nuclear probe and protein of interest channels within a bright field channel-based mask.

### In vitro T-cell activation

Primary T cells were isolated from the lymph nodes and/or spleen. The erythrocytes were lysed by using ACK buffer for 2 min at RT. For the activation of T cells, T2-Kb cells were used as antigen presenting cells. T2-Kb cells were stained in RPMI with DDAO dye (2.5µM) for 15 min at 37 °C, 5% CO_2_. Next, cells were resuspended in RPMI (10% FBS, 100 U/ml penicillin, 100 mg/ml streptomycin, 40 mg/ml gentamicin) and split into a 48-well plate and loaded with 1 µM OVA peptide for 2 h at 37 °C, 5% CO_2_. Isolated OT-I cells were co-incubated with OVA-loaded T2-Kb cells in 2:1 ratio or with T2-Kb cells not loaded at 37 °C, 5% CO_2_, or with 1 µM OVA peptide for 1 h. Subsequently the cells were processed for flow cytometry or imaging cytometry analysis.

### Infection experiments

10^4^ to 10^5^ CD8^+^ OT-I T cells in 200 μl PBS were adoptively transferred into Ly5.1 congenic host mice intravenously (i.v.). On the following day, the mice were infected with 5,000 CFU/200 µl PBS of transgenic *Listeria monocytogenes* (*LM)* expressing OVA or its lower affinity variant (Q4R7) i.v., or 31.5 PFU/30 µl PBS of transgenic Influenza A-OVA (IAV, strain A/Puerto Rico/8/34, obtained from Carolyn King, University of Basel) intranasally.

Spleen and/or lungs were isolated at indicated time points and analyzed by flow cytometry. IAV-infected mice were injected i.v. with 0.5 mg/mL of anti-CD45 FITC antibody (Cat# 103108, Biolegend, Clone 30-F11) 10 minutes before euthanasia.

For the analysis of memory cells following LM-OVA infection, samples with fewer than 20 OT-I (Ly5.2^+^) events were excluded from downstream analysis of marker expression to avoid artifacts from low event counts.

### Autoimmune diabetes

Model of autoimmune diabetes was described previously (*21, 22*). Briefly, T cells isolated from *Lck* WT or KO OT-I *Rag2*^*KO/KO*^ mice were adoptively transferred to RIP.OVA mice i.v. (10^3^ cells/200 μl). On the following day, bone marrow-derived dendritic cells were matured with 100 ng/ml LPS (Sigma-Aldrich) and loaded with 200 nM of OVA peptide (SIINFEKL) for 3 h. 10^6^ OVA-loaded BMDCs were injected i.v. The glucose in urine was monitored with test strips (GLUKOPHAN, Erba Lachema) on daily basis for 2 weeks. The blood glucose was measured using Contour blood glucose meter (Bayer) on day 7 post-immunization. The animal was considered diabetic when the concentration of glucose in urine was higher than 1000 mg/dl for 2 consecutive days.

### Histology and immunofluorescence

Pancreases were harvested from RIP.OVA mice on day 6 post-immunization. For immunofluorescence microscopy, tissue sections were fixed with 4% formaldehyde (Sigma-Aldrich, #HT501128) for 3 h and subsequently left in 30% sucrose (Sigma-Aldrich, #S9378) in PBS overnight. The following day, the tissues were mounted in OCT embedding compound (Sakura Finetek Tissue-Tek, #4583), and frozen at –80 °C. 10-µm-thick tissue sections were cut using Cryostat (Leica Microsystems, CM1950) and mounted on SuperFrost Plus Adhesion slides (Erpedia, #J1800AMNZ). Dry slides were stored at –80 °C.

For conventional light microscopy, tissue sections were fixed with acetone for 15 min and air-dried for 20 min. Staining was performed with hematoxylin M (Biognost, #HEMM-OT) for 3 min and washed with tap water for 10 sec. Following, the samples were differentiated with 0.1% hydrochloric acid (Penta chemicals, #19350) in ddH_2_O for 10⍰sec, rinsed with tap water for 6 min and stained with eosin Y (Biognost, #EOY-10-OT) for 1⍰min. Again, washed with water for 30 sec. Dehydration of samples was completed with 96% ethanol for 3⍰min, 100% ethanol for 2×5 min followed with isopropanol/96% ethanol (1:1) for 5⍰min, and xylene (Lach-ner, #20060) for 2×5⍰min. Samples were mounted in water-free solakryl medium. The pictures were acquired using Widefield Leica DM6000 microscope equipped with Leica K3C color camera and HCX PL APO 40x/0.75 DRY PH2 objective.

For the Immunofluorescence staining, samples were fixed with 4% formaldehyde for 15 min, washed with PBS for 3×5 min and blocked using 5% goat serum in PBS/0.3% Triton X-100 for 1 h. Next, samples were stained overnight at 4 °C with the Insulin primary antibody and next day washed with PBS for 3×5 min and stained with the secondary antibody goat anti-guinea pig AF488 for 1 h at RT. Then, samples were washed with PBS for 3×5 min and were stained with the mix of primary antibodies: CD4 in AF647 (Biolegend), and CD8α primary antibody for 1 h at RT, followed by washing with PBS for 3×5 min and staining with the secondary antibody goat anti-rabbit AF555 for 1 h at RT. After washing with PBS for 3×5 min, nuclei were stained with 5 µM DAPI solution (Invitrogen, D21490) for 15 min at RT. Samples were washed with PBS for 5 min and were mounted with Prolong Gold Antifade Mountant (Invitrogen, Cat# P36930). Images were acquired using confocal microscope Leica TCS SP8 (Leica Microsystems, objective HC PL APO CS2 63×/1.40 OIL).

### Proximity Ligation Assay

CD8^+^ CD5^+^ T cells from the spleen and lymph nodes of OT-I *Rag2*^*KO/KO*^ mice were sorted and frozen in cryovials in 1 ml of freezing solution (10% DMSO (Cat# D2650, Sigma) 90% FBS). Thawed 10^5^ cells were suspended in RPMI (w/o FBS and antibiotics) and placed on Diagnostic microscope slides (Thermo Scientific, ER-201B-CE24) and incubated 45 min - 1 h at 37 °C. The cells were stimulated with 5 µg/ml anti-CD3ε (Invitrogen Cat# 16-0031-85, Clone 2C11) or left unstimulated 5 min incubation at 37 °C.

Cells were fixed with 2% PFA for 15⍰min at RT, permeabilized with 0.5% saponin for 30⍰min and blocked. Blocked cells were stained according to the manufacturer’s instructions with the Duolink *In Situ*-Detektionsreagent Orange (Cat# DUO92007, Sigma). Primary goat anti-CD3ε (Cat# EB12592, Everest) and rabbit anti-FYN (Cat# 4023, Cell Signaling polyclonal) antibodies were incubated overnight at 4 °C. The day after, the cells were washed with Wash buffer A (Sigma #DUO82046-1EA) according to manufacturer’s instructions and incubated with the secondary antibodies: Duolink *In Situ* PLA Probe anti-rabbit PLUS and Duolink *In Situ* PLA Probe anti-goat MINUS for 1 h at 37 °C. Cells were further washed with Wash Buffer B (Sigma #DUO82046-1EA) according to manufacturer’s instructions.

Nuclei were stained with DAPI (SouthernBiotech #0100-20) Roth) diluted 1:1 in Mounting medium Fluoromount-G (Cat# 0100-01 SouthernBiotech). 4-6 images were taken per sample at ×60 with a confocal microscope (Nikon C2) and analyzed with BlobFinder.

### Immunoblotting

Splenocytes were used for the determination of endogenous FYN expression. 15×10^6^ cells were lysed in 1% *n*-Dodecyl-β-D-Maltoside (DDM, Thermo Scientific #89903) in lysis buffer (30 mM Tris pH 7.4, 120 mM NaCl, 2 mM KCl, 2 mM EDTA, 10% glycerol, 10 mM chloroacetamide (Sigma #C0267), 10 mM complete protease inhibitor cocktail (Roche #5056489001), and +/-PhosSTOP tablets (Roche #4906837001)), and cleared by centrifugation (21,130 xg) for 30 min at 4 °C. Proteins were denatured in 1× Laemmli sample buffer for 3 min at 92⍰°C.

Samples were subjected to immunoblotting with indicated primary antibodies and HRP-conjugated secondary antibodies.

### Gene knock-out in peripheral T cells

For the knockout of *Lck* Alt-R™ S.p. Cas9-GFP V3 (Integrated DNA Technologies, Cat# 10008100) was used. Three specific crRNAs *Lck* (GCGGACTAGATCGTGCAATC, GCTTTCGCCACGAAGTTGAA, GACCCACTGGTCACCTATGA; Integrated DNA Technologies) or a negative control crRNA (Integrated DNA Technologies, Cat# 1079138) were used.

The RNP complexes were assembled by following the manufacturer’s instructions. 1µl of 60 pmol Cas9-GFP protein was added to each sample (1.8 µl) and incubated for 20 min at RT.

2×10^6^ cells isolated from lymph nodes of Ly5.1 OT-I *Rag2*^*KO/KO*^ mice were nucleofected using 20 µl of electroporation buffer from the P4 Primary Cell 4D-Nucleofector™ X Kit S (Lonza Cat# 197191) and an 4D-Nucleofector® X Unit (Lonza) Cat# AAF-1003X, Cat# AAF-1003B, with DS 137 program. Post-nucleofection, cells were cultured in a complete IMDM medium, i.e., supplemented with 10% FBS (GIBCO), 100 U/ml penicillin (BB Pharma), 100 mg/ml streptomycin (Sigma-Aldrich), 40 mg/ml gentamicin (Sandoz), under standard conditions (5% CO2 at 37 °C) for 1 hour before their adoptive transfer to host mice or in vitro activation: 1.5×10^6^ cells/ml in complete IMDM media with 10 ng/ml PMA and 0.5 µM ionomycin for 24 hours and complete IMDM with 2 ng/ml recombinant IL-2 (Thermofisher # 212-12-20UG) for next 48h.

### Single cell transcriptomics

Splenocytes were collected on day 5 post LM-OVA-immunization. Samples were enriched for CD8^+^ cells by using Dynabeads™ Untouched™ Mouse CD8 Cells Kit (Cat#11417D, Invotrogen) according to manufacturer’s instructions. The cells were resuspended in HBSS and stained with with LIVE/DEAD near-IR dye, anti-CD8a BV421, anti-CD45.1 PE, and anti-CD45.2 APC antibodies together with TotalSeq-C Hashtag antibodies for multiplexing (BioLegend; MH1: #155861, MH2: #155863, MH3: #155865, MH5: #155869, MH6: #155871, MH7: #155873, MH8: #155875, MH9: #155877, MH10: #155879). Viable CD8α^+^, CD45.2^+^OT-I cells were FACS sorted.

The viability and concentration of cells after sort were measured using the TC20 Automated Cell Counter (1450102, Bio-Rad). The viability of the cells pre-loading was 85%. Cells were loaded onto a 10X Chromium machine (10X Genomics) aiming at the yield of 2,000 cells per sample. The sequencing libraries were prepared using Chromium Next GEM Single Cell 5’ Reagent Kits v2 with Feature Barcode technology for CRISPR Screening and Cell Surface Protein (10X Genomics, #PN-1000263, #PN-1000286, #PN-1000541, #PN-1000190, #PN-1000215, #PN-1000250) according to the protocol provided by the manufacturer (#CG000511 Rev C).

Resulting cDNA libraries were sequenced on NovaSeq 6000 (Illumina, NovaSeq Control Software 1.8.1) with the NovaSeq⍰6000 S4 Reagent Kits 300 cycles. The reads were then demultiplexed using Illumina BCL convert.

### Analysis of scRNAseq data

The generated reads were mapped using 10X Cell Ranger v7.1.0 to the *Mus musculus* reference genome provided by Ensembl (GRCm39, version 109)(*41*). The results were subsequently processed using Seurat v4.3.0 (*42*) in R v4.2.1. Mitochondrial genes, ribosomal genes, and genes matching the regular expression Tr[ab][vdj] (primarily TCR V(D)J genes) were removed. Cells expressing fewer than 200 genes were also excluded. Sample demultiplexing was performed using the HTOdemux function from the Seurat package. Hashtag doublets and cells lacking a sufficient number of hashtag reads were filtered out.

The preprocessed dataset was normalized and scaled using the default method in the Seurat package (v4.3.0). The top 750 variable genes were selected for dimensionality reduction via principal component analysis (PCA), followed by uniform manifold approximation and projection (UMAP) using 12 principal components. Clustering was performed with the Louvain algorithm implemented in Seurat. In a second, more stringent quality-control step, clusters consisting of dead or low-quality cells were removed. The final dataset contained 5,241 cells. Cell cycle effects were regressed out by specifying the parameter *vars*.*to*.*regress = c(“S*.*Score”, “G2M*.*Score”)* during the ScaleData step in the Seurat workflow.

To quantify the expression of gene signatures, we used the AddModuleScore function implemented in the Seurat package. We computed module scores for curated gene sets: Hallmark Interferon Gamma Response (MM3878) and Biocarta TCR Pathway (MM1504) from the Mouse MSigDB collection, and GSE9650 Naïve vs Eff CD8 Tcell DN, originally defined in human genes, which we converted to murine orthologs using the Ensembl BioMart database (release December 2021). The conversion was performed using the getLDS function from the biomaRt R package (v2.60.1) to map human HGNC symbols to mouse MGI symbols. Dot plots were created with DotPlot function implemented in the Seurat package. Heatmaps were created with package pheatmap (v1.0.12).

## Data and code availability

The raw data were deposited in GEO (NCBI) and are available under the accession number GSE304760. The code used to analyze the single-cell data was deposited on GitHub: <https://github.com/Lab-of-Adaptive-Immunity/LCK_Project>.

## Acknowledgement

We gratefully acknowledge Ladislav Cupak for technical assistance, Zdenek Cimburek and Matyas Sima (Flow cytometry facility, IMG) for cell sorting, Sarka Kocourkova and Michal Kolar (Laboratory of Genomics and Bioinformatics, IMG) for cDNA library preparations. VU, VR, AMM, VN, AA, VC, VN are students at the Faculty of Science Charles University in Prague. MP is a student at the University of Belgrade. We thank all our colleagues who provided us with valuable reagents and NIH Tetramer Core Facility (USA) for pMHC tetramers. Schemes in the figures were created with BioRender.com.

This project was funded by European Union’s Horizon 2020 research and innovation programme under grant agreement No. 101125695 (ERC Starting Grant ActSwiftly to OS), project National Institute of Virology and Bacteriology (Programme EXCELES, LX22NPO5103 to OS)—funded by the European Union—Next Generation EU, Charles University Grant Agency (260623 to VU), EMBO Scientific Exchange Grants (11264 to VU), and core funding provided by the Institute of Molecular Genetics of the Czech Academy of Sciences (RVO 68378050). SM is supported by the German Research Foundation (DFG) through BIOSS -EXC294 and CIBSS - EXC 2189 and MI1942/4-1 (Project ID: 501418856). Additional support was provided by the Czech Centre for Phenogenomics, financed through LM202303 by MEYS CR, and OP RDE CZ.02.1.01/0.0/0.0/16_013/0001789 (Upgrade of the Czech Centre for Phenogenomics: developing towards translation research, supported by MEYS and ESIF).

