## Supplemental Fig. for "LCK deficiency in CD8 T cells leads to reduced proliferation and increased effector T-cell formation in mice"

### Supplemental Material

#### Supplemental Figures 1-5

**LCK deficiency in CD8 T cells leads to reduced proliferation and increased effector T-cell formation in mice**

Valeria Uleri[1,2], Vojtech Racek[1], Marta Popovic[1], Anna Morales Mendez[1], Veronika Niederlova[1], Arina Andreyeva[1], Juraj Michalik[1], Nadine M. Woessner[3], Michaela Krupkova[4], Radislav Sedlacek[4], Susana Minguet[3,5], Ondrej Stepanek[1]

1 Laboratory of Adaptive Immunity, Institute of Molecular Genetics of the Czech Academy of Sciences, Prague, Czechia

2 Faculty of Science, Charles University, Prague, Czechia

3 Signalling Research Centres BIOSS and CIBSS, Faculty of Biology, University of Freiburg, Freiburg, Germany

4 Czech Centre for Phenogenomics & Laboratory of Transgenic Models of Diseases, Institute of Molecular Genetics of the Czech Academy of Sciences, Vestec, Czechia

5 Center of Chronic Immunodeficiency (CCI), University Clinics and Medical Faculty, University of Freiburg, Freiburg, Germany

#### Figure S1

**A Gating strategy for the analysis of lymph node cells in the steady-state**

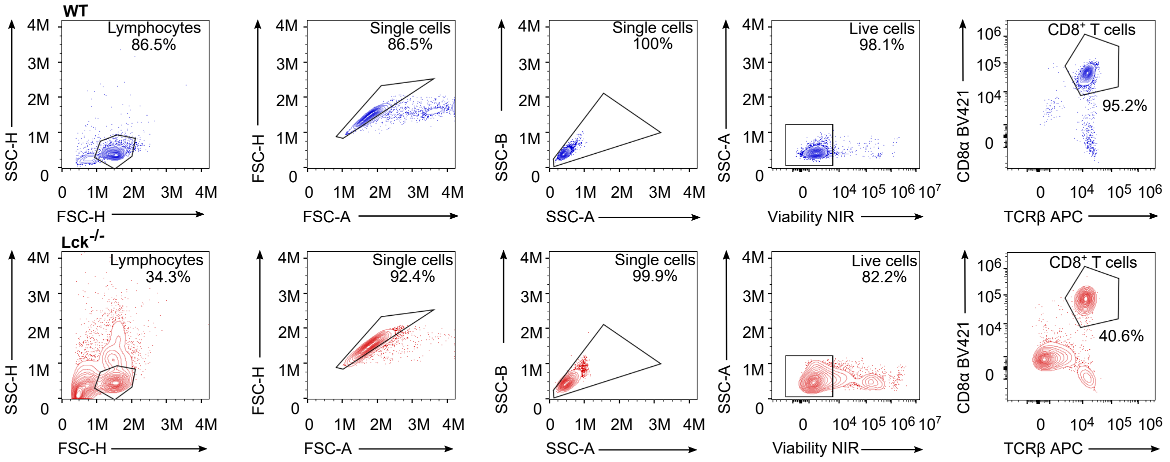


**B Phenotyping of *Lck*-/- OT-I mice**

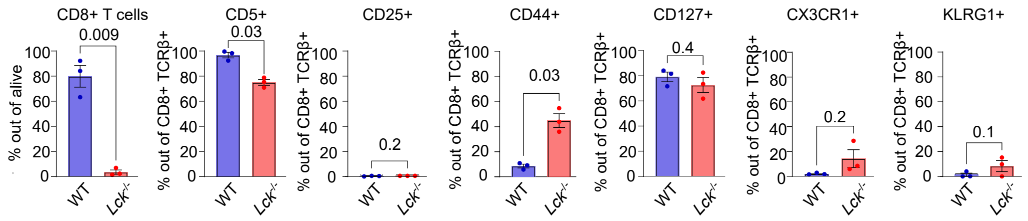


**C Gating strategy for adoptively transferred OT-I T cells**

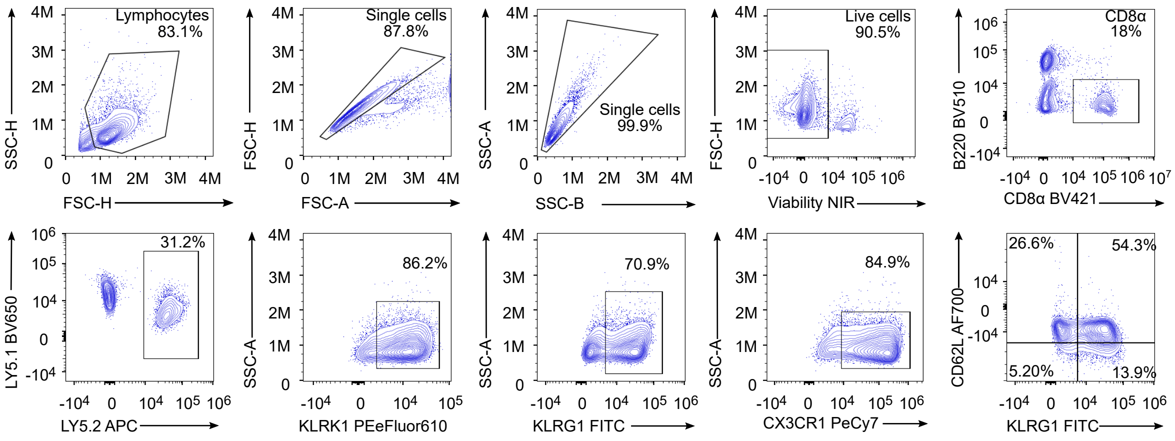

**Figure S1.**

(A) Representative gating strategy used for the analysis of live peripheral lymphocytes from lymph nodes of WT and *Lck*$^{-/-}$ OT-I *Rag2*$^{-/-}$ mice. Lymphocytes were gated based on side scatter height (SSC-H) versus forward scatter height (FSC-H), followed by FSC-H versus FSC-A and SSC-B versus SSC-A to exclude doublets. Live cells were identified by excluding Live/Dead™ Fixable Near-IR–positive events versus SSC-A plot. CD8$^+$ T cells were then gated as TCRβ$^+$CD8$^+$ double-positive cells.

(B) Mixed peripheral lymph nodes and splenocytes from WT and *Lck*$^{-/-}$ OT-I *Rag2*$^{-/-}$ mice were analyzed by flow cytometry. Mean±SEM. n = 3 mice per group in 3 independent experiments (B). Statistical significance was calculated using two-tailed paired t test.

(C) Representative gating strategy used for the analysis of live peripheral splenocytes after LM-OVA infection. Lymphocytes were first gated based on side scatter height (SSC-H) versus forward scatter height (FSC-H), followed by doublet exclusion using FSC-H versus FSC-A and SSC-B-H versus SSC-A. Live cells were identified by exclusion of Live/Dead™ Fixable Near-IR–positive events. CD8$^+$ T cells were gated as CD8$^+$ and B220$^-$ cells. Transferred OT-I cells were identified as Ly5.1$^-$ Ly5.2$^+$. OT-I T cells positive for KLRG1, KLRK1, and CX3CR1 were gated as indicated.

#### Figure S2

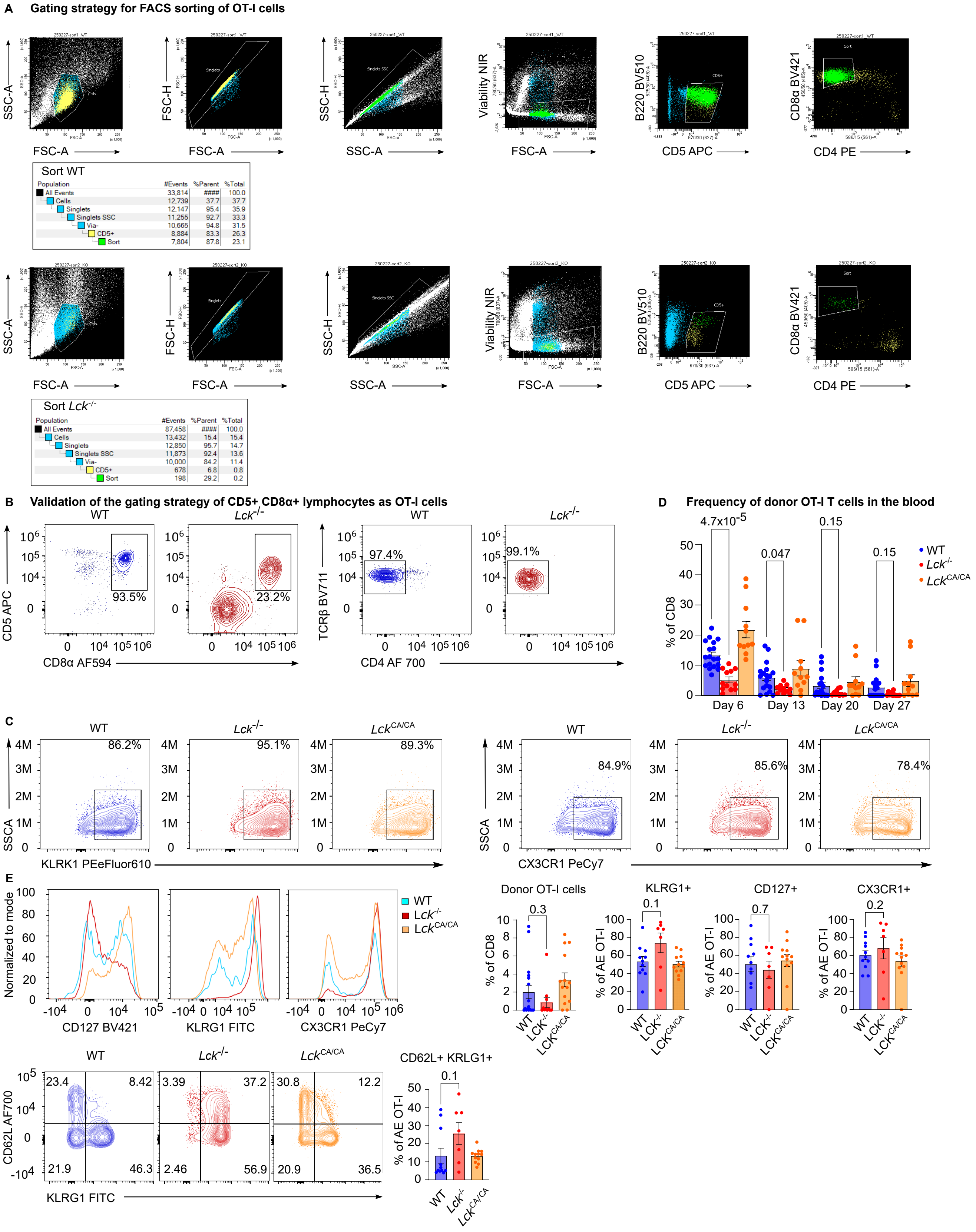


Sort WT

| Population | #Events | %Parent | %Total |
|---|---|---|---|
| All Events | 33,814 | #### | 100.0 |
| Cells | 12,739 | 37.7 | 37.7 |
| Singlets | 12,147 | 95.4 | 35.9 |
| Singlets SSC | 11,255 | 92.7 | 33.3 |
| Via- | 10,665 | 94.8 | 31.5 |
| CD5+ | 8,884 | 83.3 | 26.3 |
| Sort | 7,804 | 87.8 | 23.1 |

Sort *Lck*$^{-/-}$

| Population | #Events | %Parent | %Total |
|---|---|---|---|
| All Events | 87,458 | #### | 100.0 |
| Cells | 13,432 | 15.4 | 15.4 |
| Singlets | 12,850 | 95.7 | 14.7 |
| Singlets SSC | 11,873 | 92.4 | 13.6 |
| Via- | 10,000 | 84.2 | 11.4 |
| CD5+ | 678 | 6.8 | 0.8 |
| Sort | 198 | 29.2 | 0.2 |

**Figure S2.**

(A) Gating strategy used for FACS sorting positive $CD8^+$ $CD5^+$ cells out of peripheral lymphocytes from WT and *Lck*$^{-/-}$ OT-I *Rag2*$^{-/-}$ mice. First, lymphocytes were identified based on side scatter area (SSC-A) versus forward scatter area (FSC-A), followed by doublet exclusion using FSC-H versus FSC-A and SSC-H versus SSC-A. Viable cells were gated by excluding Live/Dead™ Fixable Near-IR–positive events. T cells were identified as $CD5^+B220^-$ cells, and $CD8^+$ T cells were further gated as $CD8^+CD4^-$.

(B) Representative contour plots of cells from lymph nodes of WT and *Lck*$^{-/-}$ OT-I *Rag2*$^{-/-}$ mice. Cells were gated first as $CD8\alpha^+$ $CD5^+$ double positive, and subsequently as $TCR\beta^+$ $CD4^-$.

(C-E) $1\times10^4$ T cells from WT, *Lck*$^{CA/CA}$, or *Lck*$^{-/-}$ OT-I *Rag2*$^{-/-}$ mice were FACS sorted as $CD8\alpha^+CD5^+$ and adoptively transferred to Ly5.1 host mice that were subsequently infected with LM-OVA. (C) Splenocytes were analyzed by flow cytometry on day six post-infection. Representative contour plots showing KLRK1, and CX3CR1 expression. n = 6 mice per group in 2 independent experiments.

(D) Blood samples were analyzed by flow cytometry on indicated days post-infection by flow cytometry. n = 18 for WT, 12 for *Lck*$^{-/-}$ , and 11 for *Lck*$^{CA/CA}$ on day 6. n = 18 for WT, 11 for *Lck*$^{-/-}$, and 11 for *Lck*$^{CA/CA}$ on day 13. n = 17 for WT, 11 for *Lck*$^{-/-}$, and 11 for *Lck*$^{CA/CA}$ on days 20 and 27 in 4 independent experiments.

(E) Splenocytes were analyzed by flow cytometry on day 30 post-infection. Representative histograms and aggregate data show the frequency of OT-I cells and expression of indicated surface markers. n = 17 for WT, 11 for *Lck*$^{-/-}$, and 14 for *Lck*$^{CA/CA}$ for determination of OT-I frequency. 6, 4, and 4 mice from WT, *Lck*$^{-/-}$, and *Lck*$^{CA/CA}$, respectively, were removed from the analysis of surface markers, because the number of detected OT-I cells was below the arbitrary limit of 20.

Mean±SEM. Statistical significance was calculated using two-tailed Mann-Whitney test.

**Figure S3**

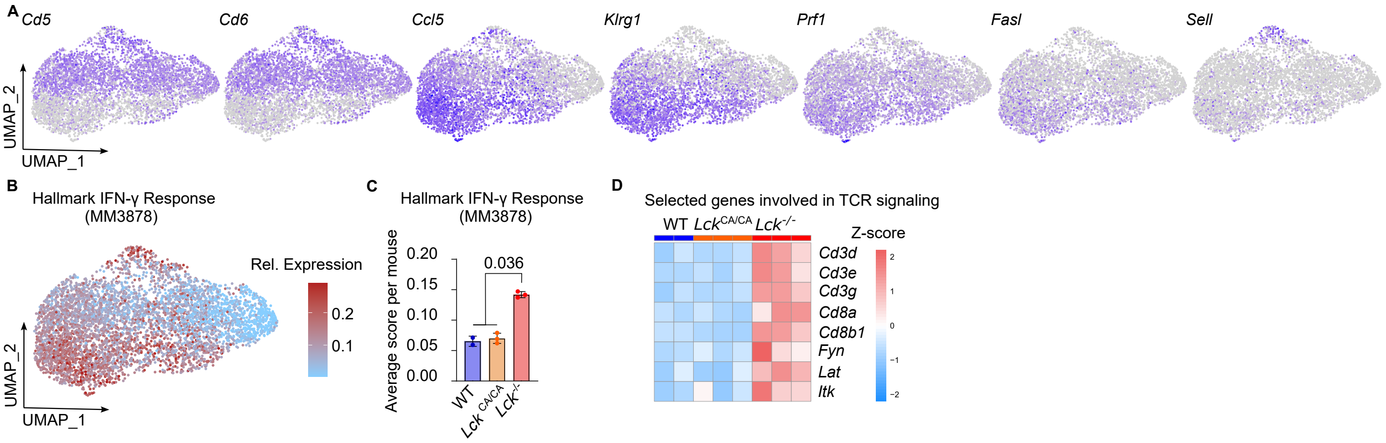
A
Cd5
Cd6
Ccl5
Klrg1
Prf1
Fasl
Sell
UMAP_2
UMAP_1
B
Hallmark IFN-γ Response (MM3878)
Rel. Expression
0.2
0.1
UMAP_2
UMAP_1
C
Hallmark IFN-γ Response (MM3878)
Average score per mouse
0.20
0.15
0.10
0.05
0.00
0.036
WT
Lck CA/CA
Lck -/-
D
Selected genes involved in TCR signaling
WT Lck CA/CA Lck -/-
Z-score
Cd3d
Cd3e
Cd3g
Cd8a
Cd8b1
Fyn
Lat
Itk

**Figure S3.**

(A-D) Additional analyses of the scRNAseq experiment shown in Fig. 2.

(A) UMAP visualization shows the log-normalized expression level of selected genes (*Cd5, Cd6, Ccl5, Klrg1, Prf1, Fasl, Sell*) in OT-I $CD8^+$ T cells as shades of blue. n = 2 mice per group for WT, n = 3 mice per group for *Lck*$^{CA/CA}$ and *Lck*$^{-/-}$.

(B) UMAP visualization shows the gene module score of Hallmark Interferon Gamma Response gene set (MM3878, mouse MSigDB collection).

(C) Average Hallmark Interferon Gamma Response (MM3878, mouse MSigDB collection) module score per mouse. WT, *Lck*$^{CA/CA}$ and *Lck*$^{-/-}$ OT-I $CD8^+$ T cells. Mean±SEM. Statistical significance between *Lck*$^{-/-}$ and WT+*Lck*$^{CA/CA}$ was calculated using two-tailed Mann-Whitney test.

(D) Heatmap showing the average expression of selected genes (*Cd3d, Cd3e, Cd3g, Cd8a, Cd8b1, Fyn, Lat, Itk*) from Biocarta TCR Pathway gene set (MM1504, mouse MSigDB collection) per mouse.

**Figure S4**

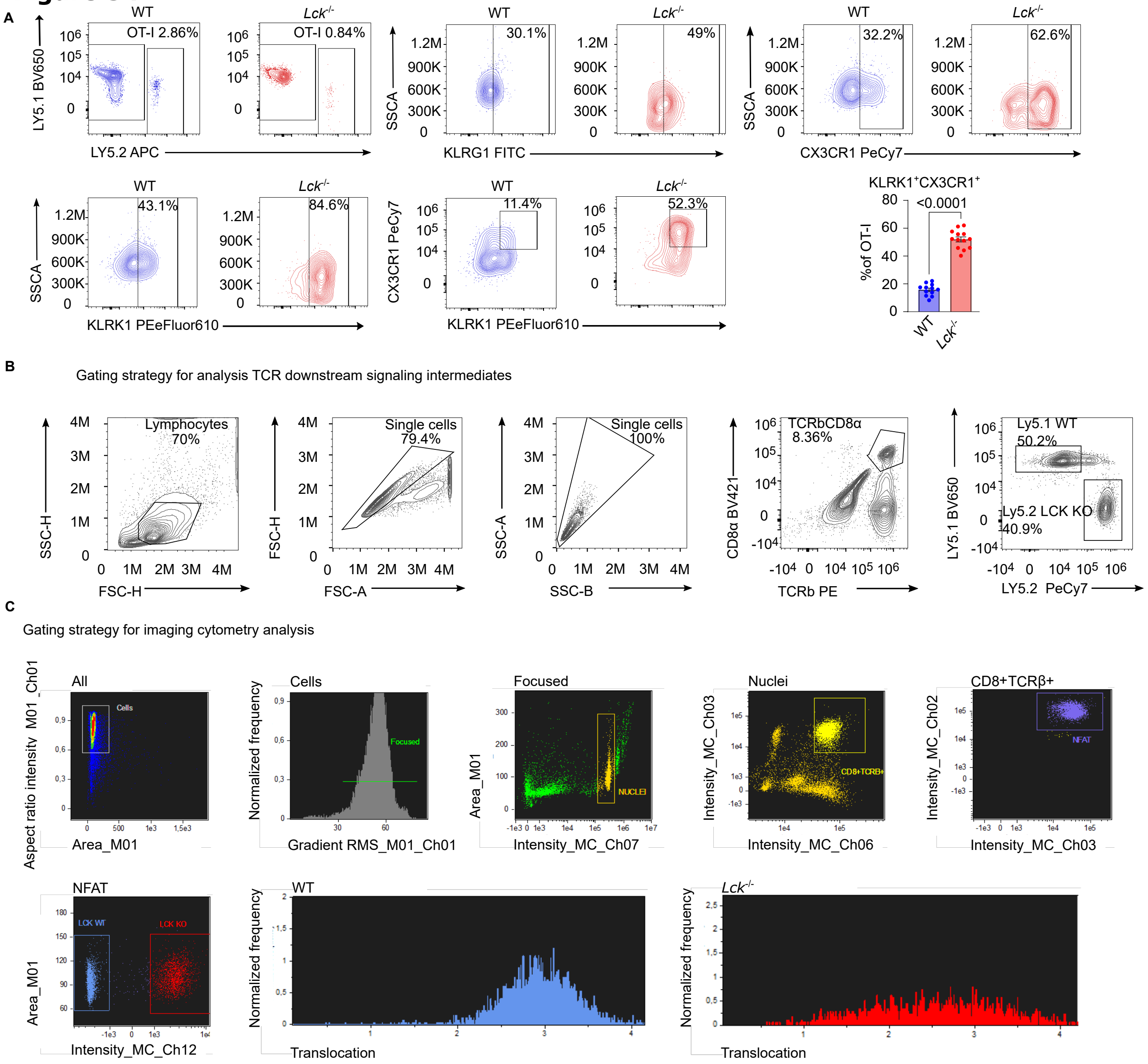
A
WT
Lck-/-
OT-I 2.86%
OT-I 0.84%
LY5.1 BV650
LY5.2 APC
30.1%
49%
SSCA
KLRG1 FITC
32.2%
62.6%
CX3CR1 PeCy7
43.1%
84.6%
KLRK1 PEeFluor610
11.4%
52.3%
KLRK1+CX3CR1+
<0.0001
% of OT-I
B
Gating strategy for analysis TCR downstream signaling intermediates
Lymphocytes 70%
Single cells 79.4%
Single cells 100%
TCRbCD8α 8.36%
Ly5.1 WT 50.2%
Ly5.2 LCK KO 40.9%
SSC-H
FSC-H
FSC-A
SSC-A
SSC-B
CD8α BV421
TCRb PE
LY5.1 BV650
LY5.2 PeCy7
C
Gating strategy for imaging cytometry analysis
All
Cells
Focused
Nuclei
CD8+TCRβ+
Aspect ratio intensity_M01_Ch01
Area_M01
Normalized frequency
Gradient RMS_M01_Ch01
Intensity_MC_Ch07
NUCLEI
Intensity_MC_Ch03
Intensity_MC_Ch06
CD8+TCRB+
Intensity_MC_Ch02
Intensity_MC_Ch03
NFAT
LCK WT
LCK KO
Intensity_MC_Ch12
Translocation

**Figure S4.**

(A) $1 \times 10^3$ OT-I T cells were FACS-sorted as CD8α$^+$ CD5$^+$ from WT, *Lck*$^{-/-}$ OT-I *Rag2*$^{-/-}$ mice and adoptively transferred to Ly5.1 host mice, which were infected with LM-OVA. The cells were analyzed on day seven post-infection. Representative plots and the quantification of KLRK1$^+$ CX3CR1$^+$ effector cells are shown. Mean±SEM. n = 12 for WT and n = 13 of *Lck*$^{-/-}$ in 3 independent experiments. Statistical significance was calculated using two-tailed Mann-Whitney test.

(B-C) Representative gating strategies used for the analysis of signaling intermediates by flow cytometry (B) and imaging cytometry (C).

(B) Lymphocytes were first gated based on SSC-H versus forward FSC-H, followed by FSC-H versus FSC-A and SSC-B versus SSC-A to exclude doublets. CD8$^+$ T cells were gated as CD8$^+$TCRβ$^+$, Ly5.1 WT and Ly5.2 *Lck*$^{-/-}$ OT-I cells were the distinguished based on the expression of Ly5.1 and Ly5.2 markers.

(C) Lymphocytes were first gated by aspect ratio intensity (Ch01) versus area (M01). Focused cells were identified based on normalized frequency versus gradient RMS (Ch01). Singlets were gated using area (M01) versus DAPI nuclear stain intensity (Ch07). CD8$^+$ T cells were identified as double-positive for TCRβ (Ch06) and CD8 (Ch03). NFAT-positive CD8$^+$ T cells were further gated based on NFAT (Ch02) versus CD8 (Ch03) intensity. Ly5.1 WT and Ly5.2 *Lck*$^{-/-}$ OT-I cells were distinguished using NFAT (Ch02) versus Ly5.2 (Ch12). Final histograms show nuclear localization score for NFAT in Ly5.1 WT and Ly5.2 *Lck*$^{-/-}$ OT-I cells.

### Figure S5

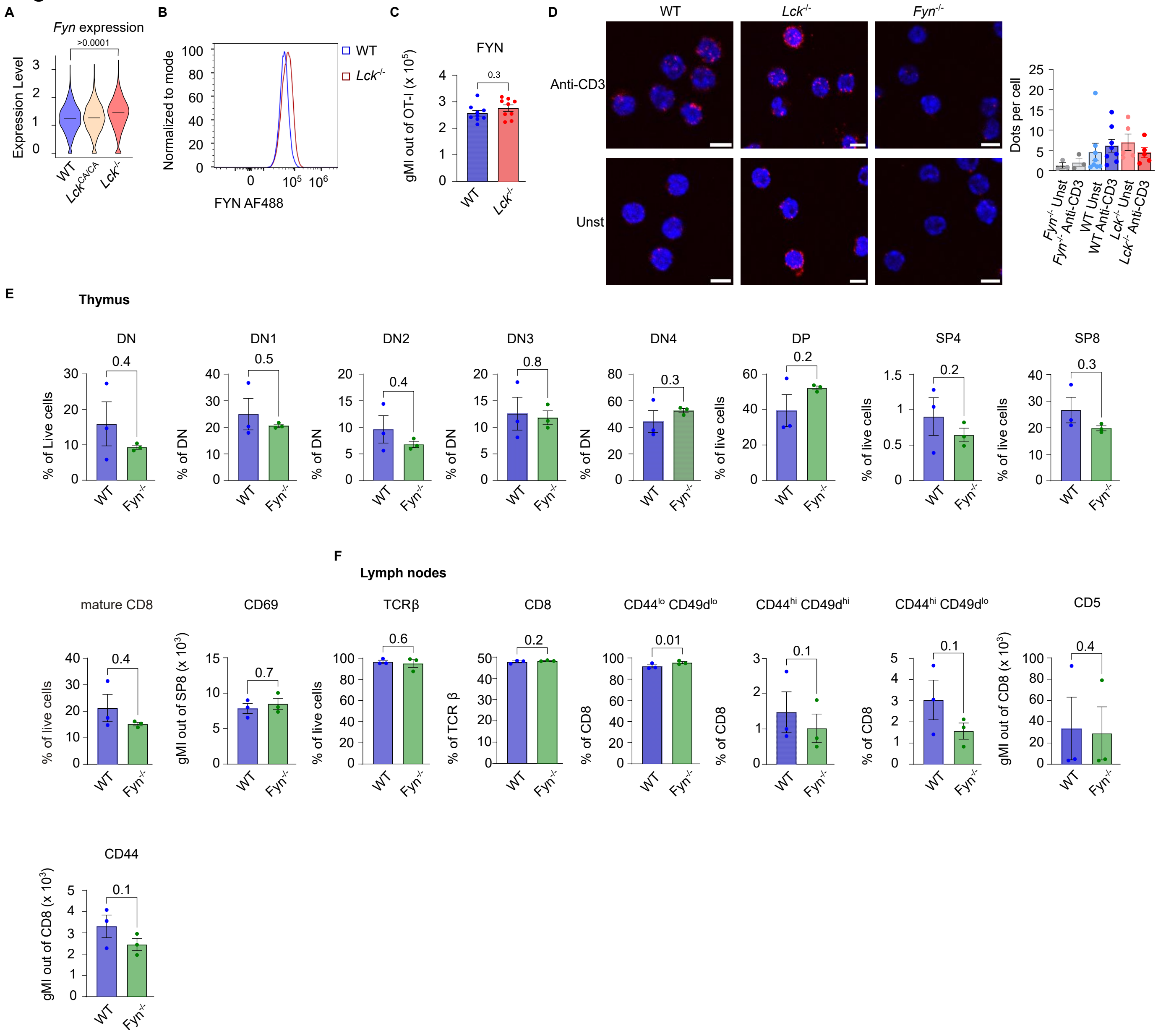
A
Fyn expression
>0.0001
Expression Level
WT
Lck CA/CA
Lck -/-
B
Normalized to mode
FYN AF488
WT
Lck -/-
C
FYN
0.3
gMI out of OT-I (x 10^5)
WT
Lck -/-
D
WT
Lck -/-
Fyn -/-
Anti-CD3
Unst
Dots per cell
Fyn -/- Unst
Fyn -/- Anti-CD3
WT Unst
WT Anti-CD3
Lck -/- Unst
Lck -/- Anti-CD3
E
Thymus
DN
DN1
DN2
DN3
DN4
DP
SP4
SP8
% of Live cells
% of DN
% of live cells
WT
Fyn -/-
F
Lymph nodes
mature CD8
CD69
TCRβ
CD8
CD44lo CD49dlo
CD44hi CD49dhi
CD44hi CD49dlo
CD5
CD44
% of live cells
gMI out of SP8 (x 10^3)
% of TCR β
% of CD8
gMI out of CD8 (x 10^3)

**Figure S5.**

(A) Log-normalized expression of *Fyn* in WT, $Lck^{CA/CA}$, and $Lck^{-/-}$ OT-I $CD8^+$ T cells responding to LM-OVA on day five post-infection in scRNAseq experiment shown in Fig. 2. Mean is indicated by the horizontal line. n = 2 mice for WT, n = 3 for $Lck^{CA/CA}$ and $Lck^{-/-}$. Statistical significance between genotypes was calculated using two-tailed Mann-Whitney test.

(B) FYN expression in lymph nodes from WT and $Lck^{-/-}$ mice measured by flow cytometry. A representative experiment out of 2 in total.

(C) $1\times10^4$ OT-I T cells were FACS-sorted as CD8α+CD5+ from WT or $Lck^{-/-}$ OT-I $Rag2^{-/-}$ mice and adoptively transferred to Ly5.1 mice that were infected with LM-OVA. Expression of FYN on day six post-infection was measured by flow cytometry. n = 8 mice for WT and n = 9 mice for $Lck^{-/-}$ in 3 independent experiments. Statistical significance was calculated using two-tailed Mann-Whitney test.

(D) In situ proximity ligation assay for detecting the proximity (<80 nm, red dots) of CD3ε and FYN in purified naïve primary T cells from WT, $Lck^{-/-}$, and $Fyn^{-/-}$ OT-I $Rag2^{-/-}$ mice, either left unstimulated (Unst) or stimulated with anti-CD3ε (145-2C11) at 37 °C for 5 min. Nuclei were stained with DAPI. Scale bars 5 μm. Representative images and quantification of dots per cell. Mean±SEM. n = 3 independent experiments for $Fyn^{-/-}$ and $Lck^{-/-}$, n = 8 independent experiments for WT mice.

(E-F) Frequency of indicated T-cell subsets in the thymus (E) and lymph nodes (F) of WT, or $Fyn^{-/-}$ OT-I $Rag2^{-/-}$ mice. Mean±SEM. n = 3 mice per group in 3 independent experiments. Statistical significance was determined using two-tailed paired t test.
